# *Trypanosoma brucei* Tim50 Plays a Critical Role in Cell Cycle Regulation and Parasite Infectivity

**DOI:** 10.1101/2021.04.26.441502

**Authors:** Anuj Tripathi, Ujjal K Singha, Ayorinde Cooley, Taneisha Gillyard, Evan Krystofiak, Siddharth Pratap, Jamaine Davis, Minu Chaudhuri

## Abstract

Tim50 is a receptor subunit of the preprotein-translocase of the mitochondrial inner membrane, TIM23. *Trypanosoma brucei*, the infective agent for African trypanosomiasis, possesses a homologue of Tim50 (TbTim50) with a pair of characteristic DXDX(T/V) phosphatase signature motifs. Here, we demonstrated that besides its protein phosphatase activity, the recombinant TbTim50 binds and hydrolyzes phosphatidic acid in a concentration-dependent manner. *In silico* structural homology models identify the putative binding interfaces that may accommodate different phospho-substrates. Interestingly, TbTim50 depletion in the bloodstream form (BF) of *T. brucei* reduced cardiolipin (CL) levels and decreased mitochondrial membrane potential (ΔΨ). TbTim50 knockdown (KD) also reduced the population of G2 phase and increased G1 phase; thus, BF cell growth was reduced. Confocal and electron microscopy revealed a defect in regulation of kinetoplast (kDNA) replication due to TbTim50 KD. Depletion of TbTim50 increased the levels of AMPK phosphorylation, and parasite morphology was changed to stumpy-like with upregulation of few stumpy marker gene expressions. Importantly, we observed that TbTim50-depleted parasites were unable to establish infection in mice and rats. Proteomics analysis showed reductions of the translation factors, flagellar transport proteins, and many proteasomal subunits, including the mitochondrial HslVU that is known to play a role in kDNA replication. Reduction of the level of HslV in TbTim50 KD cells was further validated by immunoblot analysis. Altogether, our results showed that TbTim50 is essential for mitochondrial function, regulation of kDNA replication, and cell cycle in the BF. Therefore, TbTim50 is an important target for structure-based drug design to combat African trypanosomiasis.

**Importance:** African trypanosomiasis, a neglected tropical disease caused by parasitic protozoan *Trypanosoma brucei*, is transmitted by the tsetse fly prevalent in sub-Saharan Africa. During its digenetic life cycle, *T. brucei* undergoes multiple developmental changes to adapt in different environments. *T. brucei* BF, dwelling in mammalian blood, generates ATP from glycolysis and hydrolyzes ATP in mitochondria for inner membrane potential. We found that TbTim50, a HAD-family phosphatase, is critical for *T. brucei* BF survival *in vitro* and *in vivo*. Depletion of TbTim50 in BF reduced CL levels and mitochondrial ΔΨ and caused a detrimental effect on many cellular functions. Cells accumulated in G1-S phase, and kinetoplast was over-replicated due to depletion of mitochondrial proteasomes, HslVU, a master-regulator of kDNA replication. Cell growth inhibition was accompanied by changes in morphology, AMPK phosphorylation, and upregulation of stumpy-specific gene expression. TbTim50 is essential for *T. brucei* survival and an important *T. brucei* therapeutic target.

## Introduction

*Trypanosoma brucei* is a group of parasitic protozoa and the infectious agent of a fatal disease in human and domestic animals, known as African trypanosomiasis (1). The disease is transmitted by the bite of the tsetse fly that is prevalent in sub-Saharan Africa. Within mammalian blood, *T. brucei* exists as a proliferative long-slender (LS) bloodstream form (BF) (2, 3). The LS form is covered with a thick surface coat consisting of the variant surface glycoprotein that periodically changes and protects the parasite from the host’s immune attack (4). At the peak of each parasitic wave, the LS form is differentiated to the non-dividing stumpy (ST) BF by a cell density-sensing phenomenon. The current understanding articulates that oligopeptides secreted by the LS form generates an autocrine signaling mechanism via activation of the AMP-activated protein kinase (AMPK) and increases the expression of the ST-specific genes (5, 6). The gene expression pattern of each of these developmental forms has been widely investigated in *T. brucei*, and the signaling mechanisms involved in these processes have been identified gradually (7, 8).

*T. brucei* possesses a concatenated structure of mitochondrial DNA known as kinetoplast (kDNA) that consists of thousands of mini, and few dozen of, maxi circular DNAs. The kDNA disc is attached to the flagellar basal body through the mitochondrial membrane via a filamentous structure known as the tripartite attachment complex (TAC) (9, 10). kDNA plays a crucial role during cell division (11). kDNA duplication and segregation occurs before nuclear duplication and division. Duplicated kDNA, basal body, flagella, and nucleus are separated into two daughter cells during cytokinesis along with the division of single mitochondrion (11, 12). Despite complex structure, mitochondrial DNA in *T. brucei* only encodes 18 proteins. Therefore, as similar in other eukaryotes, a vast majority of mitochondrial proteins are encoded in the nuclear genome and imported into mitochondria after synthesis in the cytosol (13, 14).

Mitochondrial protein import machinery is overall conserved among fungi and animals. Three major complexes are present: the translocase of the mitochondrial outer membrane (TOM) and two translocases of the mitochondrial inner membrane TIM23 and TIM22 (15, 16). Nuclear DNA-encoded mitochondrial proteins with either N-terminal or internal targeting signal cross the outer membrane (OM) through the TOM complex (17) and select either TIM23 or TIM22 complexes, respectively, to reach the destination (18). The core components of the TIM23 complex are Tim17, Tim23, and Tim50 (19, 20). The first two components each have four TMs that form the import channel; Tim50 has single TM with a large C-terminal domain exposed in the IMS that acts as the receptor for the preprotein (21, 22). *T. brucei* mitochondrial protein import machinery is significantly divergent. Instead of two TIM complexes, *T. brucei* likely possesses a single TIM (13, 14). The major component of the *T. brucei* TIM (TbTIM) is TbTim17 (23). Several other trypanosome-specific proteins are found associated with TbTim17, including TbTim62, TbTim42, TbTim54, two Rhomboid-like proteins, Acetyl CoA dehydrogenase, six small TbTims, and TbTim50, which is relatively conserved. Functions of these proteins have not been explored fully (23–28).

We identified the Tim50 homologue in *T. brucei* (TbTim50) and showed that it is involved in the import of preproteins into mitochondria (24). Similar to its homologues in other eukaryotes, TbTim50 possesses a characteristic C-terminal domain containing a pair of signature motifs, DXDX(T/V) (24), found in the class of CTD-phosphatases (Sc-FCP1/h-SCP1) that dephosphorylate the serine residues in the tail of the RNA polymerase II large subunit (29). The members of these CTD-phosphatases also are involved in other cellular functions, such as regulation of proteasomal activity, stress tolerance, and dephosphorylation of different signaling factors (30, 31). Altogether, these proteins belong to a large superfamily known as haloacid dehalogenase (HAD), universally present in both prokaryotes and eukaryotes (32). Substrates for HAD phosphatases vary widely and include small metabolites, proteins, and other macromolecules. Lipin, which hydrolyzes phosphatidic acid to diacyl glycerol and phosphate, also belongs to this family (32). Previously, we demonstrated that the recombinant TbTim50 possesses a dual-specific protein phosphatase activity (24). Later, we demonstrated that TbTim50 downregulation in the procyclic form (PF) of *T. brucei* that dwells in the insect vector increased tolerance to oxidative stress by increasing the levels of the phospho-tyrosyl phosphatase-interacting protein, PIP39 (33, 34). PIP39 is a similar HAD-family phosphate but localized in the glycosomes that are peroxisome-like organelles harboring primarily the glycolytic enzymes in trypanosomatids (5). The communication between these phosphatases likely is mediated via AMPK phosphorylation due to decreased production of ATP in TbTim50 KD PF (34). Here, we investigate the role of TbTim50 in the BF. We found that TbTim50 depletion is more detrimental in BF than PF. TbTim50 KD caused mitochondrial dysfunction and reduced the levels of the mitochondrial proteosomal subunit HslVU, a master regulator of kDNA replication, thus arrest cell growth both *in vitro* and *in vivo*. Therefore, TbTim50 is essential for the parasite survival in its mammalian host.

## Results

### TbTim50 can accommodate phosphoepitope binding and possesses PA-phosphatase activity

We demonstrated previously that the recombinant TbTim50 (rTbTim50) could dephosphorylate both serine/threonine and tyrosine phospho-peptides, and mutation of the conserved aspartate residues, D242 and D244, in TbTim50 abolished this activity (24). As a similar DXDX(T/V) motif was found in TbLipin (35, 36), we wanted to investigate whether TbTim50 also possesses phosphatidic acid (PA) phosphatase (PAP) activity. For this purpose, we purified the recombinant TbTim50 with a glutathione-S-transferase (GST) tag (rTbTim50-GST) at the C-terminus. The GST protein was purified in parallel to use as a control (Fig. 1A). The rTbTim50-GST hydrolyzed para-nitrophenyl phosphate (pNPP) in a dose-dependent manner as expected (Fig. 1B). When PA was used as the substrate, the rTbTim50-GST released phosphate with a similar profile (Fig. 1C), whereas rGST did not show any activity either with pNPP or PA. Together, these results showed that TbTim50 phosphatase possesses a broader substrate specificity than it was thought earlier (24). Next, we performed a lipid array to check the binding affinity of TbTim50 with different phospholipids. A piece of nitrocellulose membrane containing spots for 15 different phospholipids (100 pmole each), i.e., lysoPA, lyso phosphatidyl choline, Phosphatidyl inositol (Ptdlns), Ptdlns-3P, Ptdlns-4P, phosphatidyl choline, spingosine-1-P, Ptdlns-3,4P2, Ptdlns-3,5P2, Ptdlns-4,5P2, Ptdlns-3,4,5P3, PA, phosphatidyl serine, and a blank, were used for this assay. The rTbTim50-GST showed stronger binding affinity for PA. Ptdlns-3P and Ptdlns-4P also bound with rTbTim50-GST (Fig. 1D). In contrast, rGST showed minimal binding with these lipids. Homology modeling of TbTim50 with the crystal structure of ScTim50 using the Cn3D program (www.ncbi.nlm.nih.gov) showed the presence of a similar core with multiple α-helices and β-sheet structures. However, the two antiparallel β-strands that protruded out of the core structure of ScTim50 did not match with the corresponding region in TbTim50 (Fig. 1E). In addition, TbTim50 is distinct structurally from other Tim50 homologues due to the presence of the transmembrane helix (285-310 aa) located within the CTD PPase domain (24) (Fig. S1). To identify the putative binding pockets for the PA and pNPP substrates, we generated a homology model of the TbTim50 CTD PPase domain (228-404 aa) using the predictive modeling Phyre2 program (37). The model was validated using the SAVES server (http://services.mbi.ucla.edu/saves/ramachandran) (Fig. S1). Molecular docking analyses were conducted with both the PA and pNPP substrates using Autodock vina (Fig 1F, 1G), and theoretical binding free energies were obtained (Table S1) (38). These results further confirmed that TbTim50 is a membrane-embedded enzyme capable of binding and hydrolyzing multiple substrates. These structural and docking predictions coupled with TbTim50 possessing PAP activity provides strong evidence that TbTim50 is indeed a HAD phosphatase.

**FIG 1.**
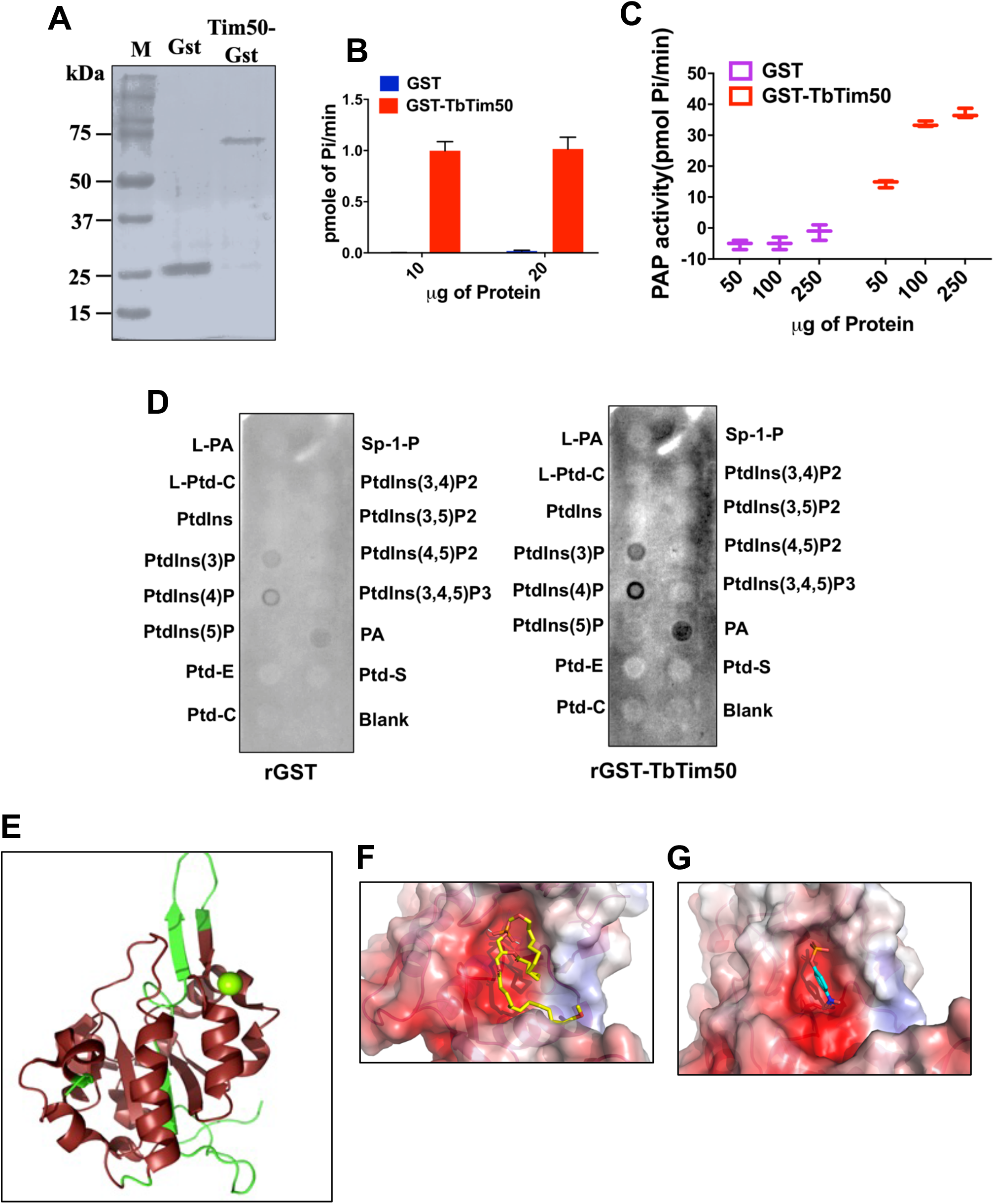
The purified recombinant TbTim50 binds and hydrolyzes PA. (A) Recombinant GST-TbTim50 (rGST-TbTim50) and GST (rGST) as control were expressed in *E.coli* and purified using affinity chromatography. (B) The phosphatase activity was measured using pNPP (0.4 M) as the substrate and varying amounts (10 and 20 μg) of purified recombinant proteins, GST-TbTim50 (red bar), or GST as control (blue bar) for 15 min. Released phosphate was quantitated by the fluorometric method as described in the material and methods. (C) Recombinant TbTim50 displays phosphatidic acid phosphatase activity. The enzymatic activity was measured by the release of phosphate from 1,2-dioctanoyl-sn-glycero-3-phosphate (DiC8 PA). The substrate was incubated with increasing amounts (50-250 μg) of either GST-TbTim50 (orange) or GST as control (purple) recombinant proteins. The amount of phosphate released was measured by using PiBlue reagent and recording the absorbance at 620 nm. Error bars in (B) and (C) were calculated from three independent experiments. (D) The rGST-TbTim50 binds with PA. PIP strip array was performed as described in the materials and methods. Spotted lipids are shown; L-PA (lyso-PA), L-Ptd-C (lyso-phosphatidyl choline), Ptdlns (phosphatidyl inositol), different Ptdlns-phosphates as indicated, Ptd-E (Ptd ethanolamine), Ptd-C (Ptd-choline), Sp-1-P (spingosine phosphate), PA (phosphatidic acid), Ptd-S (Ptd serine). (E) Structure homology modeling using the Cn3D program. The crystal structure of the ScTim50_IMS_-core region (PDB ID 4QQF) was used as the template to compare the predicted structure of TbTim50. (F) Molecular docking of phosphatidic acid (PA, ΔG= −4.8 kcal/mol) within the TbTim50 active site. (G) Molecular docking of para-nitrophenyl phosphate (PA, ΔG= −4.7 kcal/mol) within the TbTim50 active site.

### Tim50 is localized in mitochondria in the *T. brucei* BF

Previously, we reported that TbTim50 is localized in mitochondria in PF (24). TbTim50 levels are similar in the PF and ST BF, but relatively lower in LS BF (www.tritrydb.org). To confirm the sub-cellular location of TbTim50 in the BF, we *in situ* tagged TbTim50 with 12X-myc epitope at the C-terminal (due to lower affinity of the available anti-TbTim50 antibody) using the appropriate tagging vector as described in the materials and methods. Schematics for the *in situ* tagging strategy was shown in Fig. 2A. Homologous recombination of the C-terminal of the TbTim50 fragment with the endogenous locus was verified by genomic PCR analysis using gene-specific and vector-specific primers (Table S2). The gene-specific forward and reverse primers (P1 and P2) amplified the TbTim50 open reading frame (ORF) from both control and TbTim50-Myc transgenic cell lines. When the reverse primer was vector-specific (P3), expected size product was found only from transfected cell DNA (Fig. 2B). Next, we checked the expression of TbTim50-Myc by immunoblot analysis. A protein band of 75 kDa was detected by anti-myc antibody from the stable transfected cell line (Tim50-Myc) and not from the SM-BF *T. brucei* (control) (Fig. 2C). Expression of *in situ* tagged TbTim50-Myc did not have any effect on cell growth (Fig. 2D). Cell fractionation analysis showed that TbTim50-Myc is present in the mitochondrial fraction and not in the cytosolic fraction. VDAC and TbPP5 were used as the mitochondrial and cytosolic markers (Fig. 2E). Immunofluorescence microscopy revealed that TbTim50-Myc, as stained with FITC, did overlap with Mitotracker stained mitochondria in BF and PF that expressed TbTim50-Myc (Fig. 2F), whereas TbTim50-Myc stain was not overlapped with glycosomal protein aldolase (Fig. 2F). Therefore, as similar in PF, TbTim50 is localized in mitochondria in BF.

**FIG 2.**
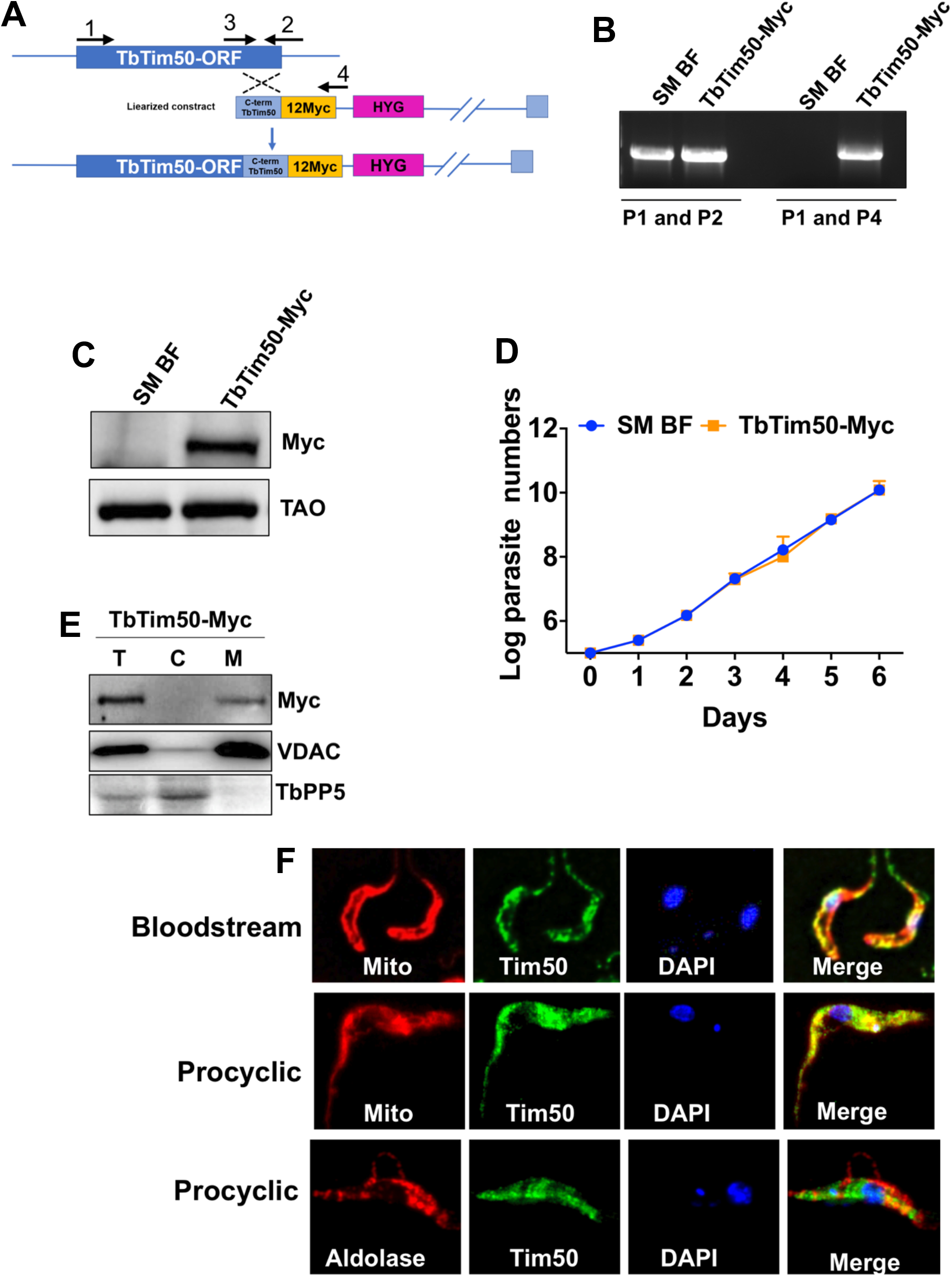
Expression and sub-cellular localization of the *in situ* Myc-tagged TbTim50 in *T. brucei*. (A) Schematic representation of the strategy used to generate *in situ* tagged TbTim50-Myc construct. After transfection, the drug-selected parasites were cloned by limiting dilution. (B) Genomic PCR analysis of the unmodified and modified locus of TbTim50 was performed to verify the integration of the construct. Location of the primers (P1–P4) used are shown in Figure 2A. The amplicons were further sequenced for confirmation. (C) Immunoblot analysis of total proteins from the parental control (SM BF) and TbTim50-Myc cell lines probed with anti-myc antibody. TAO was used as the loading control. (D) The SM BF and TbTim50-Myc cells were grown, and cell numbers were counted each day for six days. Cells were reinoculated when the parasite number reached 1×10^6^ cells/ml. The log cumulative cell number was plotted against days postinduction. Standard errors were calculated from four independent experiments. (E) Total [T], cytosolic [C], and mitochondrial [M] fractions were collected after solubilization of the cell membrane with 0.03% digitonin as described in materials and methods. Equal amounts of proteins (20 μg) were loaded per lane and immunoblotted with anti-myc, anti-VDAC, and anti-TbPP5 antibodies. VDAC and TbPP5 were used for the mitochondrial and cytosolic markers, respectively. (F) *In situ* immunofluorescence staining of TbTim50-Myc BF and PF. Live cells were stained with MitoTracker^®^ Red, fixed, and stained with anti-myc as the primary and FITC-conjugated antimouse IgG as the secondary antibodies. To localize glycosomal aldolase in the PF, anti-aldolase primary and Texas-red-conjugated anti-rabbit secondary antibodies were used. Images were taken by LSM510 confocal microscope using 60X magnification. Merged pictures are shown for colocalization.

### Tim50 knockdown in BF reduced cell growth and mitochondrial ΔΨ

In order to understand the function of TbTim50 in BF, we developed a TbTim50 RNAi cell line using the TbTim50-Myc BF as the parental control. Induction of RNAi by doxycycline showed that the level of TbTim50 transcript was reduced by 75% (Fig. 3A), and a similar reduction of the TbTim50-Myc protein level was observed (Fig. 3B). TbTim50 knockdown (KD) reduced the growth rate of the BF significantly, particularly after day four post-induction (Fig. 3C). Within 5–6 days post-induction, cell doubling time was increased about 3–4 fold. To analyze the effect of TbTim50 KD on mitochondrial ΔΨ, the control and TbTim50 RNAi cells were grown in the presence of doxycycline for two and four days and stained with MitoTracker™ Red, which is taken up by mitochondria in a ΔΨ -dependent manner (39). Cells were fixed and analyzed by flow cytometry. Results showed that TbTim50 KD reduced mitochondrial ΔΨ (~60%) by two days post-induction of RNAi (Fig. 3D & E). We did not observe any further reduction of mitochondrial ΔΨ after induction of RNAi for four days in BF. Therefore, similar to PF, demonstrated earlier, TbTim50 is required to maintain mitochondrial ΔΨ in BF. In BF, mitochondrial ΔΨ is maintained by ATP hydrolysis via the reverse action of ATP synthase. However, using immunoblot analysis we could not detect any reduction in the levels of this enzyme. As we observed that Tim50 strongly binds with PA and is capable to hydrolyze PA, a precursor for cardiolipin (CL), we wanted to examine whether TbTim50 knockdown affects the levels of CL in *T. brucei*. Cardiolipin (CL) is present predominantly in the mitochondrial inner membrane (MIM) and is essential for mitochondrial function (40). During measurement of CL levels in control and TbTim50 RNAi cells grown for four days in doxycycline-containing medium, we observed about 40% reduction of CL levels due to TbTim50 KD (Fig. 3F). This suggests that TbTim50 either is linked to CL synthesis or instability in the membrane. Loss of CL is found detrimental for mitochondrial ΔΨ in many systems. Therefore, this could be the cause of lower mitochondrial ΔΨ in TbTim50 KD BF.

**FIG 3.**
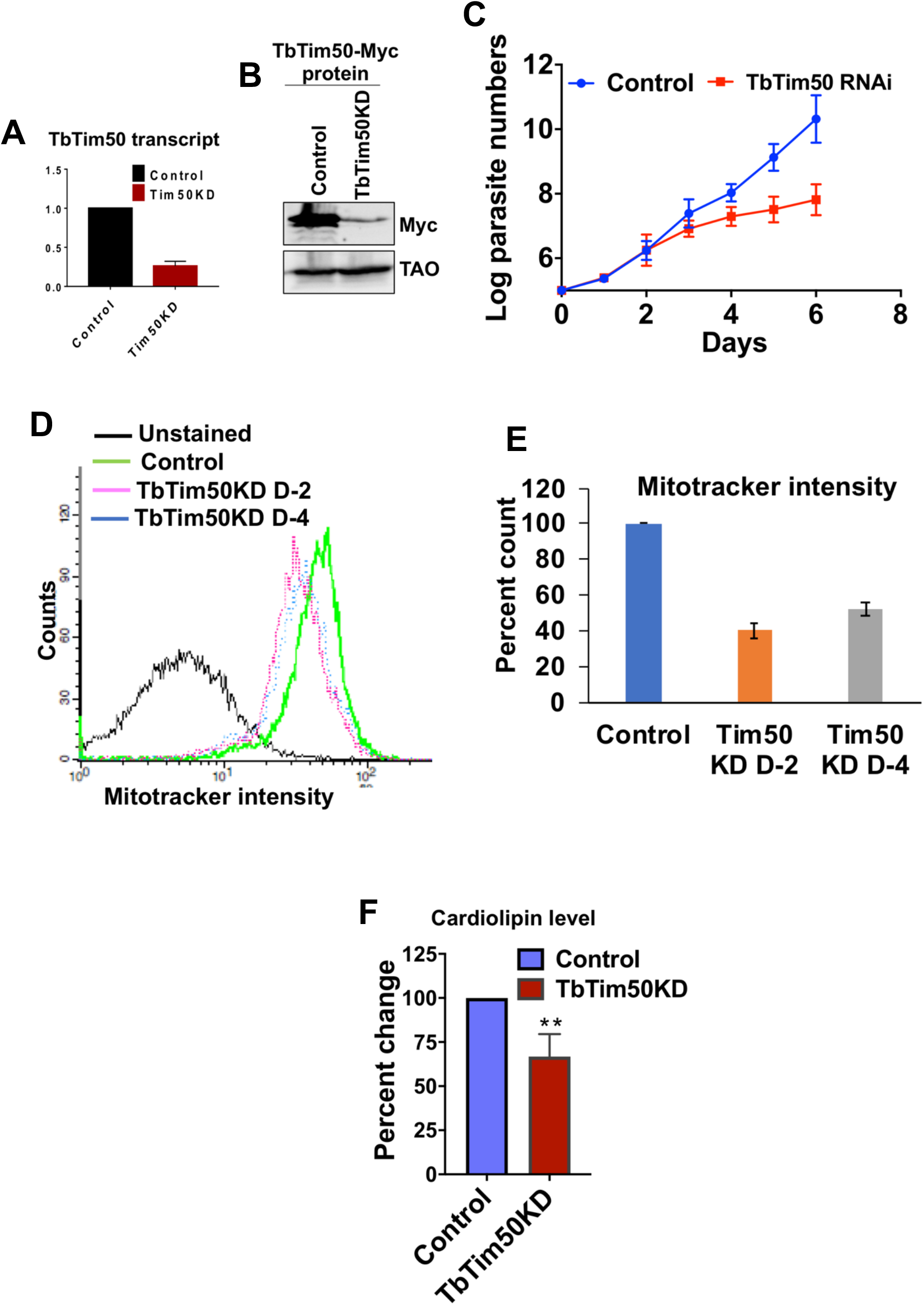
TbTim50 depletion inhibits BF cell growth, reduced cardiolipin levels, and mitochondrial ΔΨ. (A) RT-PCR analysis of the TbTim50 transcript levels in TbTim50-Myc (Control) and TbTim50-Myc/TbTim50 RNAi cells after induction of 48 h with doxycycline. The levels of the target transcript were normalized with the levels of tubulin transcript in each sample, and the normalized value for control was considered as 100%. (B) Immunoblot analysis of total proteins from TbTim50-Myc (Control) and TbTim50-Myc/TbTim50 RNAi (TbTim50 KD) cells grown in the presence of doxycycline for 48 h were probed with anti-myc antibody. TAO was used as the loading control. (C) Growth curve for the TbTim50-Myc (Control) and TbTim50-Myc/TbTim50 RNAi (TbTim50 KD) cells grown in the presence of doxycycline. Cell numbers were counted each day for six days post-induction. Cells were reinoculated when the parasite number reached 1×10^6^ cells/ml. The log cumulative cell number was plotted against days post-induction. Standard errors were calculated from three experiments. (D) Effect of TbTim50 KD on mitochondrial ΔΨ. The TbTim50-Myc (Control) and TbTim50-Myc/TbTim50 RNAi (TbTim50 KD) cells (1×10^7^) were harvested at two (D2) and four (D4) days post-induction and stained with MitoTracker^®^ Red. Fluorescence intensity was measured with a FACSCalibur (Becton Dickinson) analytical flow cytometer using absorption at 578 nm and emission at 599 nm. FlowJo software was used to analyze the results. (E) Quantitation of the fluorescence intensity from triplicate samples was performed. Fluorescence intensity of the control was considered as 100%. (F) Change in the CL-content in BF *T brucei* due to Tim50 depletion. CL levels were measured in the cell lysate of control and Tim50 KD cells as described in the materials and methods. Values shown are means ± standard errors from triplicate samples. Significance values were calculated by *t*-test and are indicated by asterisks (**, *P* < 0.01).

### Tim50 knockdown *T. brucei* could not establish infection in the host

To understand the effect of TbTim50 KD on BF cell growth *in vivo*, we infected three groups of mice with control, TbTim50 KD, and TbTim17 KD BF, respectively. We used TbTim17 KD cells to compare the effect of KD of TbTim50 and TbTim17, as both of these are Tim proteins and involved in mitochondrial protein import. We supplemented the drinking water with doxycycline to maintain the induction of RNAi *in vivo*. The group of mice infected with the control BF died within five days post-infection, as expected (Fig. 4A). The mice infected with TbTim17 KD BF survived a little longer than the control. The blood parasitemia levels were below the detection levels up to eight days; after that, the parasite number increased in the blood, and mice died within 12 days (Fig. 4A). Surprisingly, mice infected with TbTim50 KD BF did not show any symptoms; parasite levels were below 10^4^/ml of blood, and mice survived more than 3 weeks (Fig. 4A). We repeated this experiment three times and observed similar results. For further confirmation, we infected two groups of Sprague-Dawley rats with control and TbTim50 KD BF (1×10^5^ parasites/rat). The control group died within six days; the mice infected with TbTim50 KD BF survived (Fig. 4B). At day 23 postinfection, we removed doxycycline from the drinking water to reverse the RNAi effect, but we did not see any increase in the blood parasitemia levels. Next, we injected the rest of the animals with 1×10^6^ and 1×10^7^ parasites/rat at day 38 and day 52, respectively, from the initial infection day. All animals survived for 60 days. At this point, we sacrificed all rats. Parasite levels were ≤1×10^4^/ml of blood throughout the experiment, and we were unable to collect any parasites from the blood for further analysis. In culture, we observed a reduced growth rate of BF due to TbTim50 KD; however, cell growth was not ceased. It appears that the effect of TbTim50 KD is more severe on parasite growth *in vivo*. Therefore, we determine that TbTim50 is essential to establish infection by *T. brucei* BF in animals.

**FIG 4.**
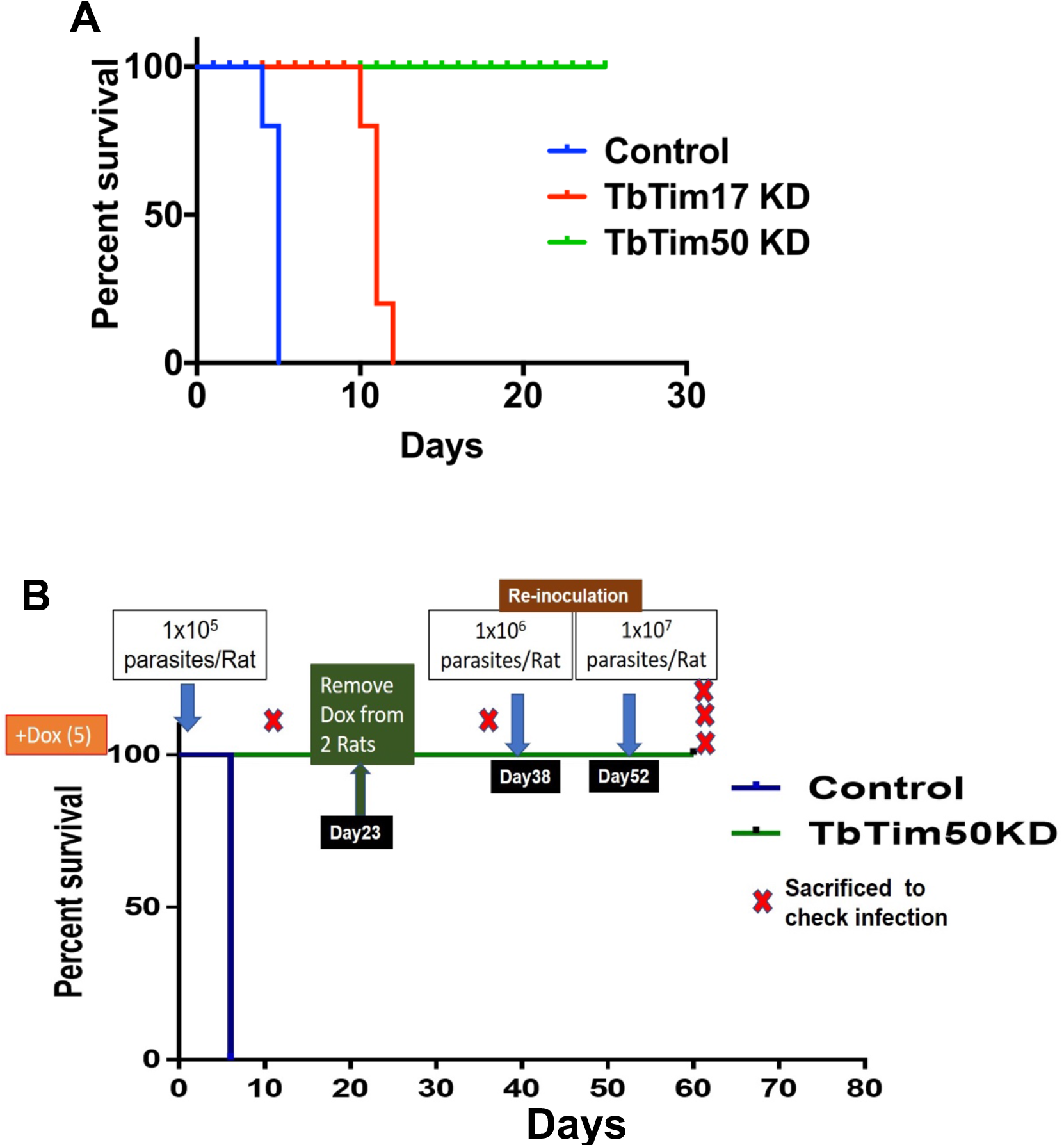
TbTim50 depletion disrupts cell cycle regulation in BF. (A) FACS profile of propidium iodide stained TbTim50-Myc/TbTim50 RNAi cells at different time points (0, 2, and 4 days) after induction of RNAi. (B) A bar graph represents the percentage of cells in each cell cycle phase (G1, S, G2) at indicated time points. (C) Fluorescence microscopy of DAPI-stained BF before and after induction of TbTim50 RNAi for 4 days. Nucleus and kinetoplast were marked as K and N, respectively. (D) Quantitation of the number of nucleus (N) and kinetoplast (K) per cell. Values shown are mean percentage of cells ± standard errors out of >200 cells of each type. Significance values were calculated by *t*-test and are indicated by asterisks (**, *P* < 0.01). (E) Electron microscopy of the control and TbTim50 KD cells at 4 days post-induction. Representative pictures are shown. Kinetoplast (K) and flagellar pocket region (F) are marked.

### TbTim50 KD hampered cell cycle regulation in BF

To elucidate the reason for *T. brucei* growth inhibition both *in vitro* and *in vivo* due to TbTim50 KD, we analyzed the cell cycle phases. For this purpose, cells were fixed with propidium iodide and the cellular DNA content was measured by FACS analysis. Results showed that TbTim50 KD decreased the population of 4C or G2-phase cells and increased the population of 2C or G1-phase cells, suggesting that transition to mitotic phase is hampered (Fig. 5A). Quantitation of the data from multiple experiments revealed that the increase and decrease of the G1- and G2-phase cell population, respectively, were about 20% (Fig. 5B), which could be accounted for inhibition of cell growth as shown in Fig. 3B and 5. During cell cycle in trypanosomatids, kinetoplast (K) division occurs first along with the basal body and flagellar duplication, followed by nuclear (N) division and cytokinesis (11,12). Therefore, at G1 phase, each cell possesses 1N1K, and during S to G2 transition, cells could have either 1N2K or 2N2K (9). By counting the numbers of N and K per cell from >200 cells each from the control and TbTim50 KD groups, we noticed significant differences in the size and numbers of N and K in TbTim50 KD cells. Compared to control, TbTim50 KD cells had larger K and N, and often K and N were segregated asymmetrically (Fig. 5C and Fig. S2); we noticed that the proportion of 2N2K cells decreased significantly (Fig. 5D). In contrast, 1N large-K and 1N with unsegregated-K were accumulated. These results showed TbTim50 KD cells are defective in kDNA replication/segregation; S-phase prolonged, thus created larger N and K. Analysis of *T. brucei* ultrastructure using electron microscopy (EM) revealed that the kDNAs appear more electron dense and longer on average in TbTim50 RNAi cells compared to control (Fig. 6). The population of the electron dense kDNA was increased 3-4 folds in TbTim50 KD BF than control. This further indicates that replication was continued without segregation of kDNA.

**FIG 5.**
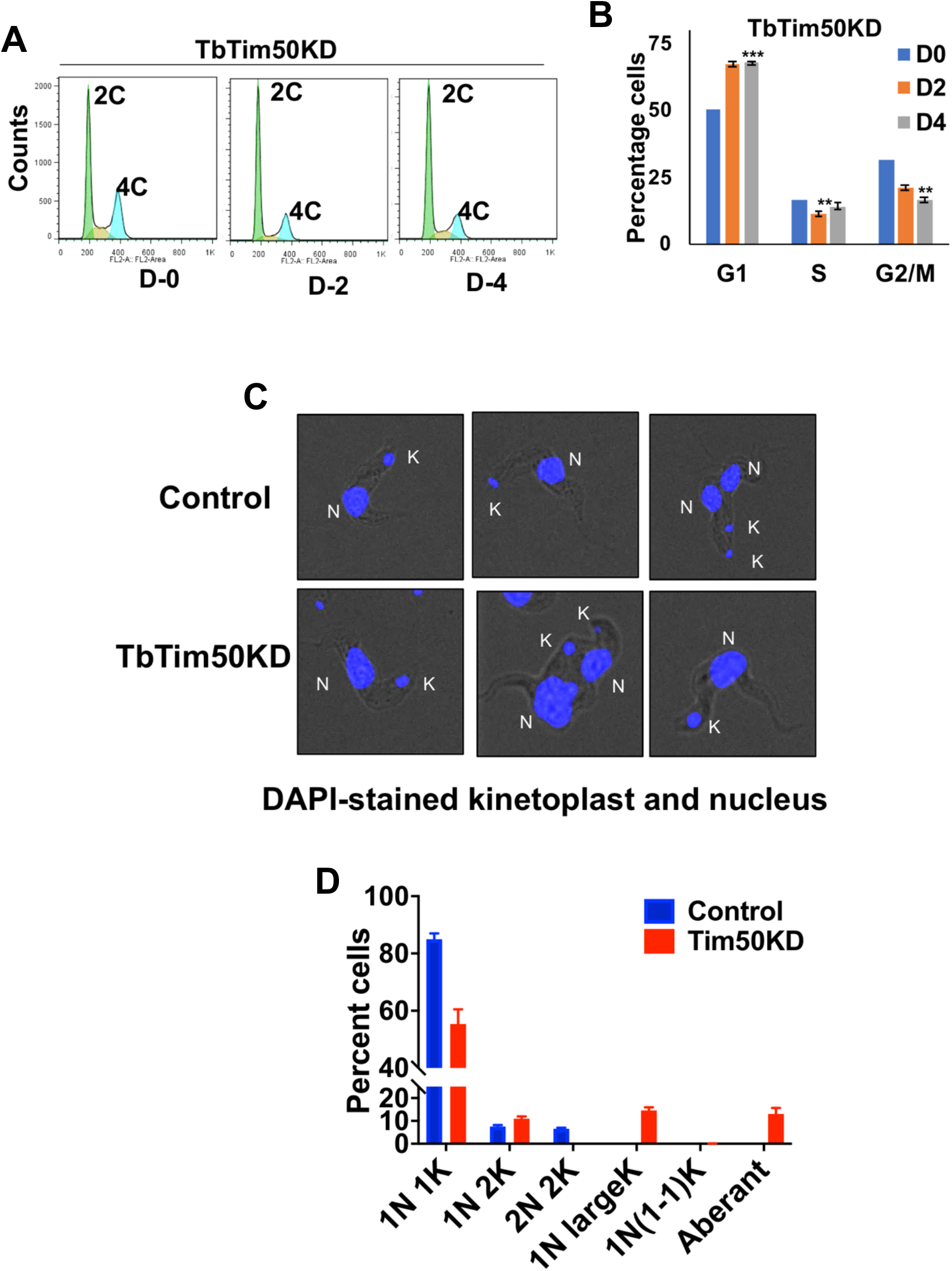
TbTim50 KD altered cell morphology, increased AMPK phosphorylation, and upregulated expression of the ST-specific transcripts. (A) Giemsa-stained TbTim50 RNAi cells at days 0, 2, and 4 after induction with doxycycline. (B) Quantitative RT-PCR analysis of the EP1, PAD1, PAD2, PIP39, and TbTim50 transcripts in control and TbTim50 RNAi *T. brucei* BF cells grown for 2 days in the presence of doxycycline. Telomerase reverse transcriptase was used as the internal control. Relative levels of the marker transcripts compared to the parental control were plotted. Averages and standard errors were calculated from three experiments. (C) Immunoblot analysis of total proteins from the control and TbTim50 KD cells harvested after 4 days post-induction using different antibodies against PIP39, phospho-AMPK, PGK, and TAO as probes. (D) Densitometric analysis the protein bands for PIP39_upper (PIP39_U), PIP39_lower (Pip30_L), AMPKα1P, AMPKα2P, PGK_B, and PGK_C after normalization with that for TAO. Values shown are mean fold increase ± standard error in TbTim50 KD in comparison to control from triplicate experiments. Significance values were calculated by t test and indicated by asterisks as follows; ** P< 0.05, *** P < 0.01.

**FIG 6.**
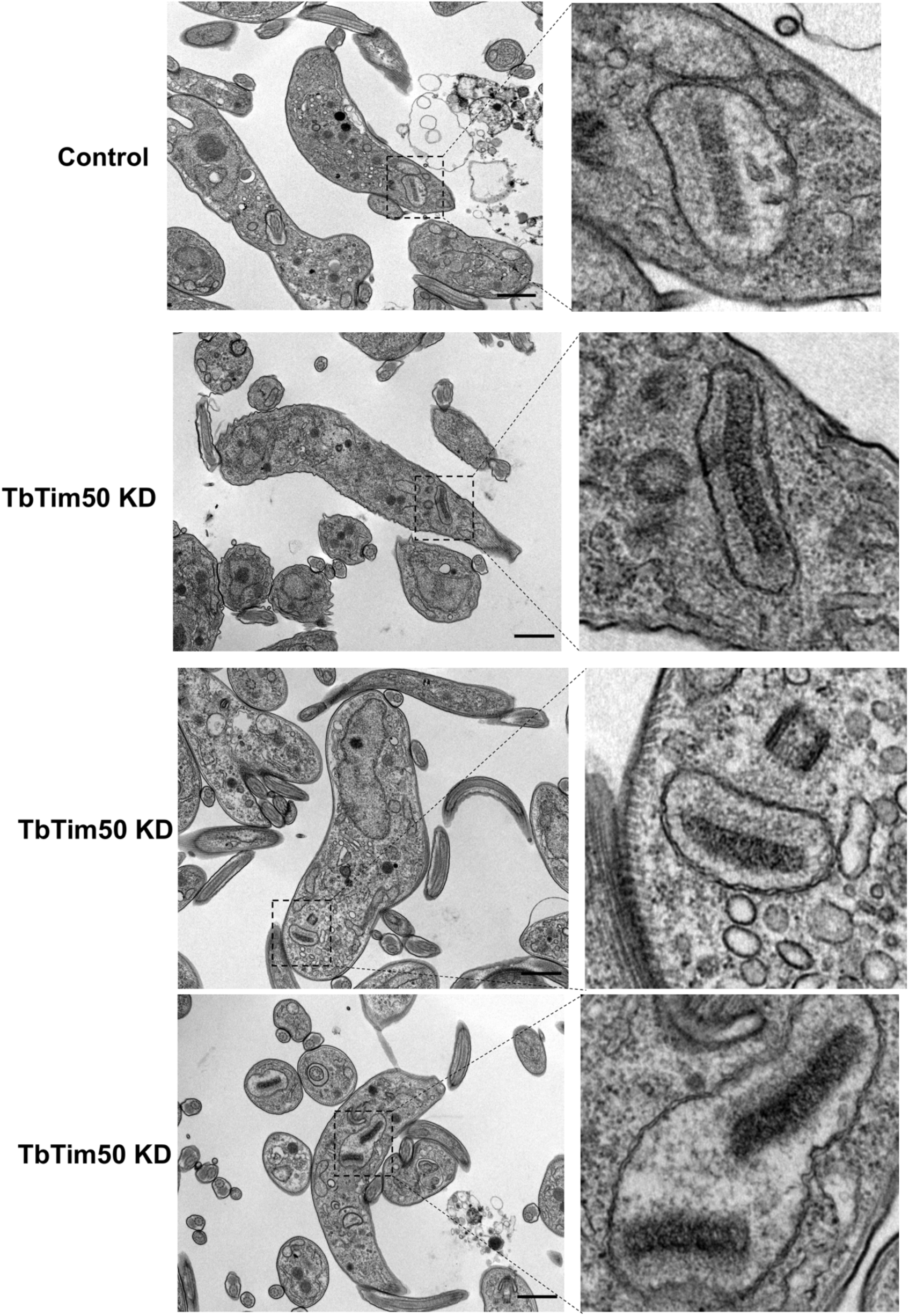
TbTim50 KD *T. brucei* is unable to establish infection in animals. (A) A survival plot of three groups of mice (five mice/group) infected with the control, TbTim17 RNAi (TbTim17 KD), or TbTim50 RNAi (TbTim50 KD) BF cells grown for 4 days in the presence of doxycycline. Mice were fed with doxycycline-containing water to continue the RNAi effect. In order to reduce the pain and distress, mice were sacrificed when blood parasitemia levels reached >2×10^8^/ml, and death was considered 24 h after. (B) Survival plot of two groups of rats (five mice/group) infected with the control and TbTim50 RNAi (TbTim50 KD) cells (1×10^5^ cells/rat) grown for 4 days in the presence of doxycycline. Rats were fed with doxycycline-containing water to continue the RNAi effect. Blood parasitemia levels were counted each day starting 2 days post-infection. At day 23, doxycycline-containing water was replaced by normal water for two experimental rats to stop the induction of RNAi. At 38 and 52 days post-infection, two rats from the experimental group were re-infected with 1×10^6^ and 1×10^7^ TbTim50 KD BF, respectively. All rats in the experimental group survived throughout the experiment. The time of euthanasia is indicated by red crosses.

### TbTim50-depleted cells have altered morphology, increased AMPK phosphorylation and increased expression levels of stumpy-specific transcripts

Examining Giemsa-stained cells under 100X objective, we found that the LS BF cells gradually appeared thicker and stumpy after induction of TbTim50 RNAi (Fig. 7A). This cell line was derived from SM427 *T. brucei* BF, which is a laboratory-adapted monomorphic cell line that loses the capacity to transform to ST form with increasing cell density (5, 6). We observed that TbTim50 KD induced these cells to stumpy-like appearance. To examine this further, we performed RT-PCR analysis for several stumpy-specific transcripts. Interestingly, we noticed several-fold upregulation of the transcript levels for PIP39, EP1 procyclin, and proteins associated with differentiation (PAD1 and PAD2) in cells where TbTim50 transcript levels were reduced by RNAi (Fig. 7B). This result further suggests that TbTim50 KD inhibits cell division and transformed these cells toward ST-like form. It has been shown that monomorphic 427 *T. brucei* could be transformed to ST-like form, when incubated with AMP or cell permeable AMP-analogue and transformation of the pleomorphic SL to ST BF by SIF is mediated via AMPK phosphorylation (5, 6). Therefore, we investigated whether AMPK phosphorylation is altered due to TbTim50 KD. Immunoblot analysis indeed showed that phosphorylated AMPK, both α1 and α2, levels upregulated due to TbTim50 KD (Fig. 5C). In contrast, the levels of PGK and TAO were unchanged. Previously, we showed that TbTim50 KD in PF increased AMPK phosphorylation and increased the levels of PIP39 protein several-fold (34). Although we observed a significant upregulation of AMPK phosphorylation and an upregulation of PIP39 transcript levels in the TbTim50 KD BF, PIP39 protein levels were increased minimally. PIP39 antibody recognized a pair of bands of expected sizes. The upper band could be the phosphorylated PIP39. The anomaly between PIP39 transcript and protein levels is likely because PIP39 protein expression is regulated further either at the stage of protein translation or stability in the monomorphic strain. Overall, we observed that TbTim50 KD disrupts cell cycle regulation possibly by inhibition of kDNA-division and induced a stage transition in BF.

**FIG 7.**
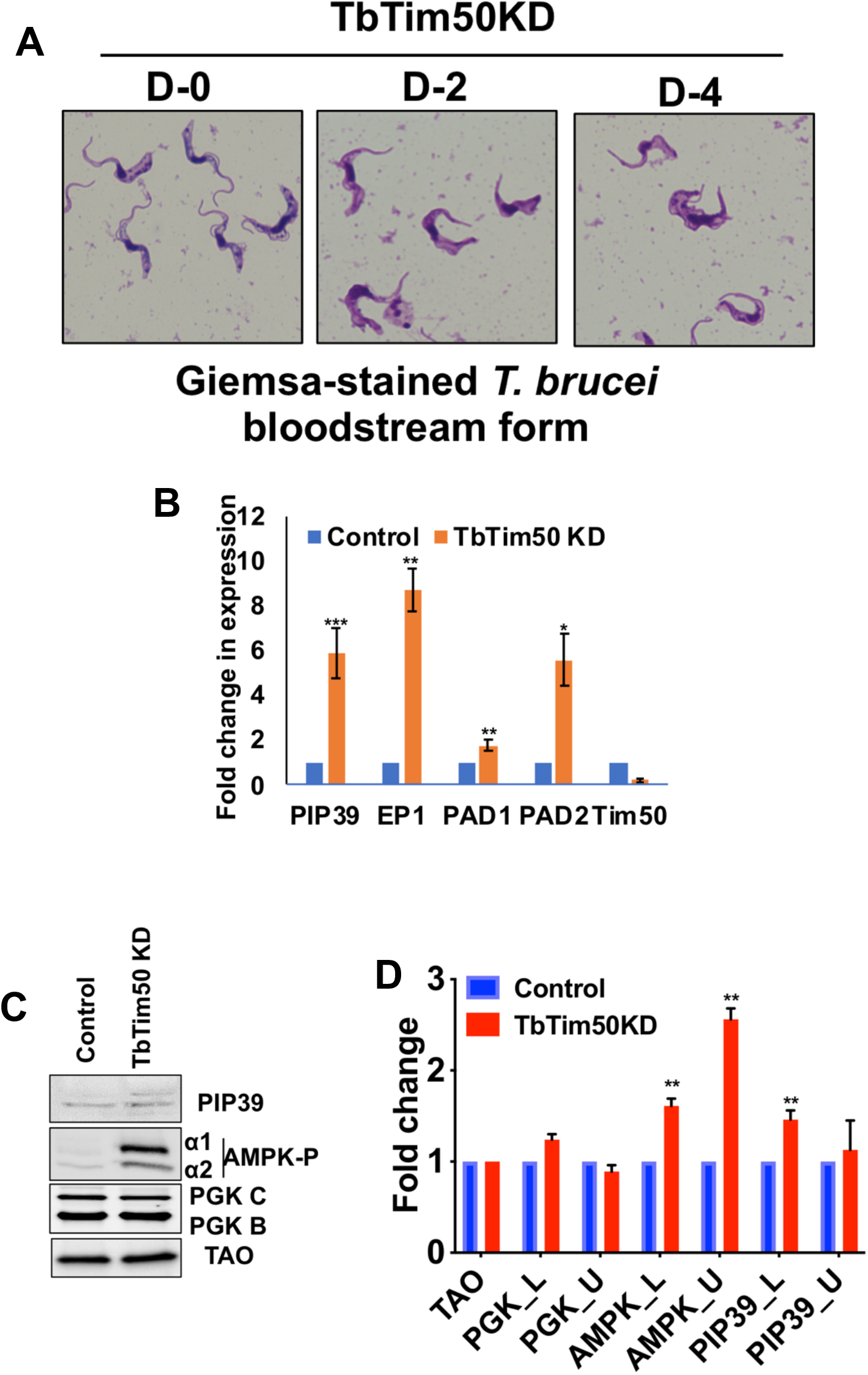
Comparative proteomics analyses and validation of the proteomic results. The semiquantitative proteomics analysis was performed to compare the proteomes in the control and TbTim50 KD cells. Statistically significant down- and upregulated proteins from three biological replicates were used for STRING analysis as described in the materials and methods. (A and B) Interaction map of proteins, those significantly downregulated (A) and upregulated (B) due to TbTim50 KD. Clusters of downregulated proteins were encircled and labeled according to the GO-term. Upregulated proteins were less clustered, and individual proteins were labelled as identified. (C) *In situ* tagged TbHslV-HA is expressed and localized in mitochondria. Immunoblot analysis of the total [T], cytosolic [C], and mitochondrial [M] fractions from TbHslV-HA cells were probed with anti-HA, VDAC, TbTim17, and TbPP5 antibodies. (D) The TbHslV-HA cells were further transfected with the TbTim50 RNAi construct to develop a stable TbHslV-HA/TbTim50 KD cell line. Immunoblot analysis of total proteins from the parental control, TbHslV-HA, and TbHslV-HA/TbTim50 KD cells, grown in the presence of doxycycline for 2 and 4 days using anti-HA antibody. TAO was used as the loading control. The black arrow indicates the specific band recognized by anti-HA antibody. (E) The working model of TbTim50 function in BF. TbTim50 is critical to maintain CL levels in mitochondria. Thus, depletion of TbTim50 hampered mitochondrial ΔΨ, created an energy crisis, and increased AMPK phosphorylation that inhibits cell growth and triggers some differentiation processes. TbTim50 also is required for TbHslVU functions to regulate the kDNA replication and cell cycle progression. The non-dividing ST-like cells generated due to TbTim50 KD are cleared rapidly from the bloodstream in a mammalian host. Therefore, the TbTim50-depleted parasite could not establish infection.

### Global proteomics analysis revealed downregulation of certain cellular functions due to TbTim50 KD

In order to understand the effect of TbTim50 KD on total cellular proteomes, semi-quantitative proteomics analyses were performed using control and TbTim50 RNAi cells grown for 2 and 4 days in the presence of doxycycline. Overall, we found more proteins were downregulated (<0.5-fold) than upregulated (>1.5-fold) in TbTim50 KD cells. From the day-2 sample, we observed 130 proteins were downregulated, and 51 were upregulated (supplemental file S1). Similarly, from the day-4 sample, 57 were up- and 377 were downregulated (supplemental file S2). After overlapping these results, we found 58 proteins were common in both samples that were altered due to TbTim50 KD (Fig. S4A). Clustering these proteins according to the functional terms revealed the presence of ribosomal (27%), translational factors (12%), RNA-binding (12%), cytoskeletal (10%), proteasomal (9%), chaperones (8%), developmental regulation (7%), metabolic (5%), and stress-regulated (3%) proteins (Fig. S4B). Next, we separated the list of up- and downregulated proteins and performed STRING analysis (Fig. 8A). From these analyses, we found that downregulated proteins were clustered into different groups that include: 1) ribosomal proteins and translation factors; 2) flagellar transport proteins; 3) RNA-binding proteins; 4) proteasome subunits; 5) metabolic enzymes; and 6) ATP-synthase subunits. Downregulation of many ribosomal proteins and translational factors indicated protein synthesis was inhibited, which is not unexpected, as we see the AMPK phosphorylation was increased due to TbTim50 KD. It is known that activated AMPK stimulates the catabolic processes and inhibits the anabolic processes like protein synthesis (41). Furthermore, we found down regulation of several proteasome subunits that are known to be linked with cell cycle regulation in *T. brucei* (42). Particularly, the mitochondrial heat shock locus ATPase (HslVU) complex, known to play roles in kDNA replication/segregation (43, 44), were downregulated >4-fold (pvalue <0.005) in TbTim50 KD cells (Table 1). We found that several subunits of the ATP synthase (complex V) complex were downregulated, which could be due to loss of CL or vice versa. Alteration of the levels of the flagellar transport proteins and metabolic enzymes are indicative also of cellular stress/adaptation due to TbTim50 KD. In contrast to downregulated proteins, upregulated proteins were less clustered (Fig. 8B). These include Ca-signaling proteins, such as calmodulin, calpain, PP2c, and PP1. Several metabolic enzymes, such as glucose-6-phosphate dehydrogenase, malate dehydrogenase, alanine amino transferase, and glycerol uptake protein were increased. Upregulation of AMP-deaminase, adenylosuccinate synthase, nucleotide phosphotransferase was indicative of the cellular demand for nucleotides, likely for longer S phase. In addition, we noticed some upregulation of several ESAG-related proteins and prostaglandin synthase. Overall, the information gathered from proteomics analyses were correlated with the phenotype of the TbTim50 KD *T. brucei*.

**Figure 8.**
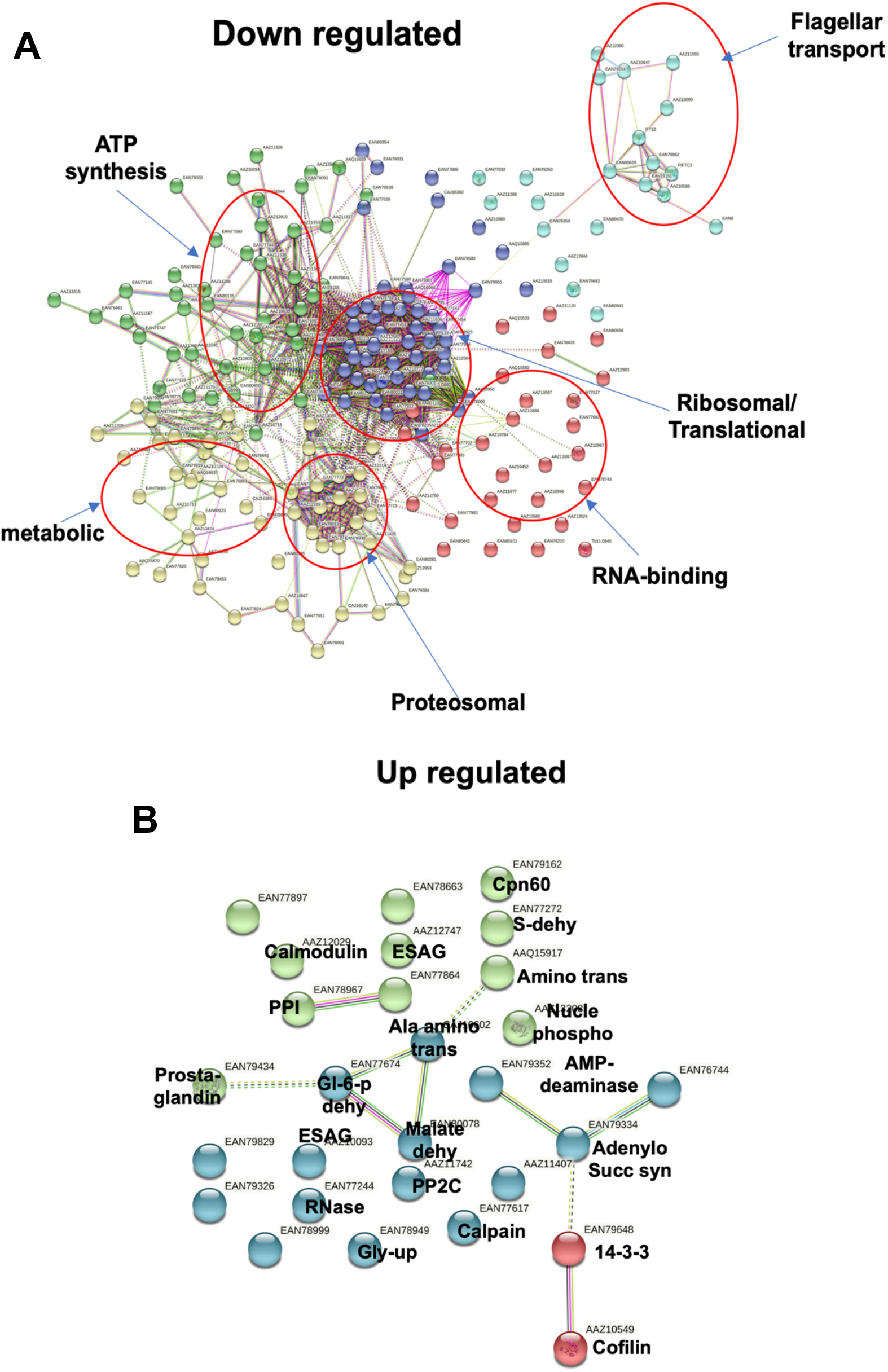

**Figure 9.**
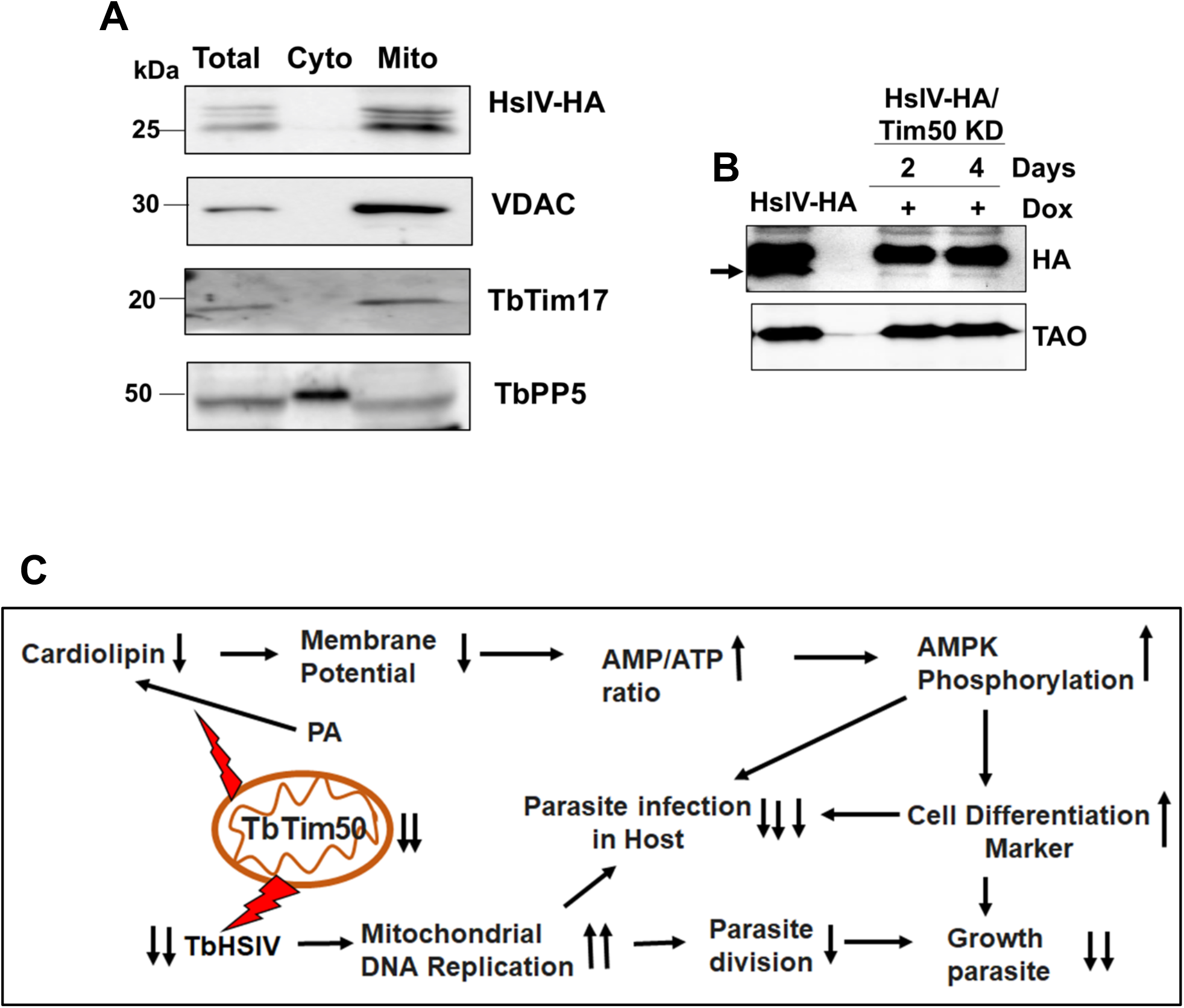

### TbTim50 depletion reduced the levels of the mitochondrial HslVU

From our proteomic analysis we observed that mitochondrial HsIVU subunits were consistently downregulated >4-fold in all biological replicates (Table 1). HslVU is a prokaryotic proteasome complex found in mitochondria in certain eukaryotic species, including trypanosomatids (43–46). *T. brucei* and related parasites thus have both mitochondrial HslVU and cytosolic 26S proteasome complexes. Interestingly, the HslVU complex has not been found in animal or human mitochondria. The HslVU complex consists of the catalytic subunit HslV and the regulatory subunit HslU (43, 45). The HslV subunit forms a dodecameric homo-oligomer, shaped like double donuts stacked on top of each other. The holo-regions on both sides are blocked by homo-hexameric HslU. HslU binds substrate proteins, unfolds them by ATP hydrolysis, and transfers to the catalytic core for degradation. *T. brucei* has two HslU subunits, U1 and U2; TbHslU1 is only able to activate TbHslV (45). Multiple reports showed that TbHslVU plays a role in kDNA replication and segregation during the cell cycle in *T. brucei* (43–46). TbHslV KD caused *T. brucei* to have larger K and N and asymmetrically divided kDNA, similar to that we found in TbTim50 KD cells. It has been shown that PIF2 helicase, which is needed for maxi circle replication, is degraded by HslVU to regulate kDNA replication and segregation (46). Therefore, we selected TbHslV to further validate our proteomics results. For this purpose, we *in situ* tagged TbHslV at the C-terminal with 3X-HA. The selected cell line showed expression of the expected size HA-tagged protein in the mitochondrial fraction (Fig. 8C). These cells were further transfected with the TbTim50 RNAi construct to observe the effect of TbTim50 KD on the levels of TbHslV. We indeed found that the TbHslV levels were reduced significantly due to depletion of TbTim50 (Fig. 8D). Therefore, our results demonstrated that reduction of the mitochondrial proteasome subunit levels is linked with the phenotype seen in TbTim50-depleted parasites.

Overall, our finding reveals that TbTim50 has two-pronged function in mitochondria. It is involved to maintain CL levels in the MIM. Thus, depletion of TbTim50 reduced mitochondrial ΔΨ and cellular ATP levels as found in CL synthase KD in BF (47). Cellular energy crisis activates AMPK by phosphorylation and induced a differentiation from SL to ST-like form (Fig. 8E). This form is rapidly cleared by host immunity and could not establish infection. TbTim50 is either directly or indirectly involved in regulation of HslVU; therefore, TbTim50 KD caused an unregulated replication of kDNA and lingers in the G1-S phase and hampered cell division (Fig. 8E).

## Discussion

Here, we investigated the role of TbTim50 in the *T. brucei* BF. Similar in PF, TbTim50 is required to maintain mitochondrial ΔΨ. TbTim50 is also critical for BF survival both *in vitro* and *in vivo*.Maintenance of mitochondrial ΔΨ is a conserved function for Tim50 among different species (48, 49). In yeast, it has been shown that Tim50 interacts with Tim23 and with the MIM via CL interaction to close the TIM23 channel in the absence of preproteins (50), thus maintaining the MIM permeability barrier. Our current understanding indicates that TbTim50 interacts weakly with TbTim17, the counterpart of the ScTim23 in *T. brucei* (14). Therefore, it is not clear whether that interaction is required to maintain mitochondrial ΔΨ. However, we found that CL levels are reduced significantly due to TbTim50 KD. As CL, the specialized lipid in the MIM plays a critical role in membrane integrity, it could be conceivable that the loss of CL is also a cause of reduced mitochondrial ΔΨ. In BF, mitochondrial ΔΨ is primarily maintained by ATP synthase, acting in reverse orientation (51). Although we did not notice any reduction of ATP-synthase α or β subunits by immunoblot analysis, semi-quantitative proteomics analysis showed reductions in the levels of α, β, and γ subunits of the ATPase complex (File S1 and S2). Loss of CL may hamper the ATPase assembly and function in the mitochondria, which could be an additive effect for leaky MIM in TbTim50 KD BF. It has been shown recently that the KD of CL synthase (CLs) in *T. brucei* BF reduced cellular ATP levels and mitochondrial ΔΨ; as a result, cell growth was reduced (47). We found a similar situation in TbTim50 KD BF due to reduction of CL.

CL is synthesized in mitochondria from PA and cytidine triphosphate (CTP) in a multi-step process (52, 53). The last step for CL synthesis in *T. brucei* has been shown to be similar to that in bacteria rather to that in fungi and mammals (54, 55). *T. brucei* CL-synthase (TbCLs) utilizes two molecules of phosphatidylglycerol (PG) to make CL, whereas CLs in other eukaryotes utilize PG and CDP-diacylglycerol (CDP-DAG). Interestingly, dephosphorylation of PGP by a phosphatase generates PG. Human phosphotyrosyl phosphate localizes to mitochondria (PTPMT1) and yeast GEP4 acts as the PGP phosphatase (57, 58). In fact, GEP4 is a member of the HAD phosphatase family (58). Therefore, involvement of TbTim50, which is also a HAD phosphatase, in CL synthesis is highly probable. Furthermore, CL stability primarily depends on its association with membrane proteins (58). Therefore, TbTim50 loss may increase CL degradation. CL oxidation also increases its degradation (59). In this regard, we have shown that TbTim50 KD increased ROS production (34), therefore it may also contribute to CL degradation. Whether TbTim50 participates in CL synthesis or the loss of TbTim50 binding with CL in TbTim50 KD parasites reduced stability of CL requires further investigation.

Other than the effect on mitochondrial ΔΨ, we observed that TbTim50 KD increased the levels of AMPK phosphorylation in BF, suggesting an energy crisis due to TbTim50 KD. In BF, ATP is not synthesized in mitochondria; instead, it is produced solely by glycolysis (60). Mitochondrial activities, such as reoxidation of the reducing equivalents generated during glycolysis via mitochondrial glycerol-3-phosphate dehydrogenase (GPDH) and trypanosome alternative oxidase (TAO), are indeed linked for a continuous flow of glucose oxidation to produce sufficient energy (61, 62). Although we did not observe any significant reduction in the steady state levels of TAO in our immunoblot analysis, we found GPDH in the list of downregulated proteins identified by proteomics analysis. Therefore, it may be estimated that the loss of mitochondrial membrane integrity hampered the electron transfer processes and slowed down glycolysis; thus, ATP levels were reduced. A similar situation has been reported for TbCLs KD (47). In addition, we found upregulation of the glycerol uptake protein and a number of amino acid metabolic enzymes in TbTim50 KD cells by proteomics analysis, which also could be an attempt for the cellular adaptation process.

Most interestingly, we found that TbTim50 KD deregulates the cell cycle controls in the BF. Cells were accumulated in the G1-S phase; thus, the proportion of cells at the G2 phase was reduced significantly due to depletion of TbTim50 levels in BF. It is known that treatment of the LS BF cells with AMP or cell permeable AMP-analogue activates AMPK by phosphorylation that triggers cells to differentiate into non-dividing ST form (5, 6). The laboratory-adapted monomorphic strains loss their capacity to transform to ST BF. Upon treatment with AMP or AMP-analogues, it appears as a ST-like morphology (63). Interestingly, we noticed that TbTim50 KD changes the monomorphic LS BF to achieve ST-like appearance. These changes were associated also with upregulation of ST-specific transcripts, including PIP39, PAD1, PAD2, and EP procyclin. Although PIP39 transcript levels were upregulated 5-fold, PIP39 protein levels were minimally increased in this cell line due to TbTim50 KD and possibly due to additional blockage at the level of translation or protein stability. Together these results showed that sudden drop in cellular energy due to mitochondrial disfunction in TbTim50-depleted cells triggered a differentiation signal, however, it was not likely completed due to a certain roadblock in the monomorphic LS BF.

Comparative proteomics analyses of the control and TbTim50 KD cells and further validation by biochemical analysis helped us to clarify why the LS BF cells were arrested at the G1-S phase in the cell cycle. Network analysis of the downregulated proteins due to TbTim50 depletion identified a cluster of proteasome subunits. These include mitochondrial HslVU complex subunits, cytosolic 26S proteasome subunits, i.e., PA26, subunit α, and non-ATPase subunits 1, 2, 3, 4, 5, 9, and 11. Both mitochondrial and cytosolic proteasomes are known to play roles in regulation of the cell cycle in *T. brucei* (42). The HslVU complex, which is unique to mitochondria in trypanosomatids and not found in its mammalian host, is of particular interest (43–46). We clearly see a downregulation of TbHslV due to TbTim50 KD not only by proteomics analysis, but also experimentally. In addition, we observed TbTim50 KD caused a similar phenotypic defect in kDNA replication/segregation as reported due to KD of the TbHslVU (44, 46). Therefore, we conclude that reduction in the levels of TbHslVU complex due to TbTim50 depletion is the cause for cell cycle dysregulation and reduced cell growth. It is likely that the reduced levels of HslVU lingered the replication and abnormal segregation of the kDNA. Kinetoplast division and segregation is the preceding steps for the downstream events such as mitosis and cytokinesis in *T. brucei*. Therefore, defects in kDNA segregation due to a reduction in the levels of HslVU may relay the signal to cytosolic proteasomal subunits that are responsible to regulate the levels of specific cyclins and other cell cycle regulatory proteins. Therefore, TbTim50 KD caused larger K and N and inhibition of cell division. As HslVU requires ATP for its function, it could be imagined that loss of energy in TbTim50 KD cells may cause this complex to become non-functional and unstable. However, reduction of HslVU was not reported due to TbCLs KD that produces a similar energy crisis (47). Therefore, it is possible that TbTim50 has a direct role in regulation of the TbHslVU complex, which could be by post-translational modification, i.e., phosphorylation/dephosphorylation. Recent data on human and yeast showed that Tim50 is the substrate for mitochondrial phosphatase Pptc7, and the phosphorylated form of Tim50 has higher activity for mitochondrial protein import (64). The phospho-proteomics datasets (Tritrypdb) showed that TbTim50 is phosphorylated at the Thr72 residue and, therefore, has the potential to be regulated by kinases/phosphatases.

Overall, we showed that TbTim50 is critical for the function of the mitochondrion in *T. brucei* BF and is essential for the parasite survival. It is crucial to elucidate how TbTim50 is linked to maintain the levels of CL, and TbHslVU in mitochondria to sustain BF cell growth both *in vitro* and *in vivo*.

## Materials and Methods

### *T brucei* cell culture and *T. brucei* transgenic cell lines

The BF single marker (SM) cell line of *Trypanosoma brucei* 427 was cultured in HMI-9 medium supplemented with 10% fetal bovine serum and 2.5 μg/ml G418 at 37°C in a CO2-incubator (5% saturation) (65). Transfection of SM BF with different constructs was performed using an Amaxa Nucleofector Kit (Lonza, Cologne, Germany) as described (TrypsRU Home).

### Phosphatidic acid phosphatase (PAP) activity assay

PAP activity was measured in a reaction mixture contained 50 mM Tris–HCl buffer (pH 7.5), 1 mM MgCl2, and 0.4 mM 1,2-dioctanoyl-snglycero-3-phosphate (DiC8 PA) (Avanti Polar Lipids) in a total volume of 50 μl. Reactions were carried out in triplicate by the addition of recombinant proteins (50–250 ng) at 30°C for 30 min. The reaction was terminated by the addition of 100 μl of PiBlue reagent (BioAssay Systems), and the color was allowed to develop at room temperature for 30 minutes. The absorbance was measured with a spectrophotometer (BioRad) at 620 nm. The amount of phosphate produced was quantified from a standard curve using potassium phosphate and the above reagent. The enzymatic activity was expressed as the number of pmol of phosphate released per minute.

### Lipid binding Assay

Phosphoinositide binding assay was performed using PIP strip (Thermofisher) that contain 100 picomoles spots of various phosphoinositides on a nitrocellulose membrane. The membrane was blocked with 3% fatty acid–free BSA in TBST buffer (50 mM Tris-HCl [pH 7.4], 150 mM sodium chloride, 0.1% Tween 20) for 1 h, followed by overnight incubation with 0.5 μg/ml of purified recombinant proteins GST-TbTim50 in blocking solution at 4°C. Purified GST were used in parallel as control. The membrane was washed three times in TBST buffer and probed with anti-GST antibody for 2 h at 25°C. Subsequently, the membrane was washed with TBST buffer three times and incubated with horseradish peroxidase (HRP)-conjugated anti-Rabbit IgG, and signal was detected using chemiluminescence.

### Measurements of Cardiolipin levels

Total cardiolipin (CL) in cell was measured by Fluorometric assay kit (Abcam) according to the manufacturer’s instructions, in a 96-well fluorescence black microtiter plate in triplicate. TbTim50 RNAi and control cells were grown in the presence and absence of doxycycline for 4 days, respectively. Both types of cells (1 x 10^7^) were harvested by centrifugation at 1500 x g for 10 minutes and wash with cold PBS twice. Cell pellets were resuspended in CL Assay Buffer and a detergent free lysis of cells were carried out by sonication in ice. Cell lysates were centrifuged at 10,000 x g for 10 minutes at 4°C and the supernatants were transferred to fresh microfuge tube. Soluble supernatant equivalent to 50 ug of proteins were used for the assay. Reactions were carried out in a 96-well plate after addition of 50 μL of the Probe-Mix to each well in a total volume of 100 ul. Blank reactions for each sample were performed in parallel without addition of the Probe-Mix. Plate was incubated in dark at room temperature for 5-10 minutes. The absorbance was measured at Ex/Em 340/480 nm using a fluorescence microplate reader (BIOtech). A standard curve was generated using calculated amount of Cardiolipin supplied in the kit. Cardiolipin levels in the cell lysates were assessed after subtraction of the blank reading from each of the sample.

#### Animal experiments

All experimental procedures were carried out in the Animal Care Facility at Meharry Medical College, Nashville, Tennessee, according to the approved protocol by the Institutional Animal Care and Use Committee. Female Balb/C mice (4-6 weeks old) were purchased (Envigo) and kept in cages (1-4 mice/cage and 1-2 rats/cage) on a 12-h daylight cycle at room temperature. Infection was initiated by the intraperitoneal (i.p.) injection of *T. brucei* BF (10^6^ cells/kg body weight). The drinking water for both the control and experimental groups was supplemented with doxycycline (1%) and sucrose (5%), 7 days prior to infection. The parasitemia level in infected animals was monitored by parasite count in blood collected by tail sniping on every alternate day. Animal health was monitored frequently during infection. Animals were euthanized if they were clinically moribund (hunched back, ruffled fur coat, hind limb paralysis, weakness) or had a parasite count of ≥5×10^8^/ml of blood.

### Others

Crude mitochondria isolation, membrane-potential measurement, cell cycle analysis, qRT-PCR, immunoblot analysis, immunofluorescence imaging, Expression and purification of the recombinant protein, electron microscopy, mass spectrometry, bioinformatics, homology modeling and molecular docking, densitometry, and statistical analysis methods are included in the supplemental material.

## Acknowledgements

This work was supported by the NIH grant 1RO1AI125662 (Chaudhuri). The Core Facilities at Meharry Medical College is supported by NIH grant U54RR026140/U54MD007593. We thank George Cross for *T. brucei* 427 and SM BF cell lines, Sam Alsford for the pNAT vectors, Keith Gull for the p2T7^Ti^-177 RNAi (phleomycin resistance) vector, and Keith Matthews for PIP39 antibody. The Proteomics and Electron Microscopy Cores at Vanderbilt University Medical Center are funded by NIH grants P30DK058404 and P30CA068485. We thank the Meharry Office for Scientific Editing and Publications for scientific editing support (S21MD000104).

## References

1. Sternberg JM, MacLean L. 2010. A spectrum of disease in Human African trypanosomiasis: The host and parasite genetics of virulence. Parasitology 137:2007–2015.

2. Quintana JF, Zoltner M, Field MC. 2020. Evolving Differentiation in African Trypanosomes. Trends Parasitol. Elsevier Ltd.

3. Silvester E, McWilliam K, Matthews K. 2017. The Cytological Events and Molecular Control of Life Cycle Development of Trypanosoma brucei in the Mammalian Bloodstream. Pathogens 6:29.

4. Mugnier MR, Stebbins CE, Papavasiliou FN. 2016. Masters of Disguise: Antigenic Variation and the VSG Coat in Trypanosoma brucei. PLoS Pathog. Public Library of Science.

5. Szöor B, Ruberto I, Burchmore R, Matthews KR. 2010. A novel phosphatase cascade regulates differentiation in Trypanosoma brucei via a glycosomal signaling pathway. Genes Dev 24:1306–1316.

6. Saldivia M, Ceballos-Pérez G, Bart JM, Navarro M. 2016. The AMPKα1 Pathway Positively Regulates the Developmental Transition from Proliferation to Quiescence in Trypanosoma brucei. Cell Rep 17:660–670.

7. Cestari I, Stuart K. 2020. The phosphoinositide regulatory network in Trypanosoma brucei: Implications for cell-wide regulation in eukaryotes. PLoS Negl Trop Dis 14:e0008689.

8. Cayla M, McDonald L, Macgregor P, Matthews KR. 2020. An atypical DYRK kinase connects quorum-sensing with posttranscriptional gene regulation in Trypanosoma brucei. Elife 9:1–52.

9. Mensa-Wilmot K, Hoffman B, Wiedeman J, Sullenberger C, Sharma A. 2019. Kinetoplast Division Factors in a Trypanosome. Trends Parasitol. Elsevier Ltd.

10. Schneider A, Ochsenreiter T. 2018. Failure is not an option - mitochondrial genome segregation in trypanosomes. J Cell Sci 131.

11. Gluenz E, Povelones ML, Englund PT, Gull K. 2011. The Kinetoplast Duplication Cycle in Trypanosoma brucei Is Orchestrated by Cytoskeleton-Mediated Cell Morphogenesis. Mol Cell Biol 31:1012–1021.

12. A Ploubidou, D R Robinson, R C Docherty, E O Ogbadoyi KG. 1999. Evidence for novel cell cycle checkpoints in trypanosomes: kinetoplast segregation and cytokinesis in the absence of mitosis - PubMed. J Cell Sci.

13. Harsman A, Schneider A. 2017. Mitochondrial protein import in trypanosomes: Expect the unexpected. Traffic 18:96–109.

14. Chaudhuri M, Darden C, Gonzalez FS, Singha UK, Quinones L, Tripathi A. 2020. Tim17 updates: A comprehensive review of an ancient mitochondrial protein translocator. Biomolecules. MDPI AG.

15. Neupert W, Herrmann JM. 2007. Translocation of proteins into mitochondria. Annu Rev Biochem. Annu Rev Biochem.

16. Schmidt O, Pfanner N, Meisinger C. 2010. Mitochondrial protein import: From proteomics to functional mechanisms. Nat Rev Mol Cell Biol. Nature Publishing Group.

17. Drwesh L, Rapaport D. 2020. Biogenesis pathways of α-helical mitochondrial outer membrane proteins. Biol Chem. De Gruyter.

18. Bauer MF, Sirrenberg C, Neupert W, Brunner M. 1996. Role of Tim23 as voltage sensor and presequence receptor in protein import into mitochondria. Cell 87:33–41.

19. Rehling P, Wiedemann N, Pfanner N, Truscott KN. 2001. The mitochondrial import machinery for preproteins. Crit Rev Biochem Mol Biol. CRC Press LLC.

20. Rassow J, Dekker PJT, Van Wilpe S, Meijer M, Soll J. 1999. The preprotein translocase of the mitochondrial inner membrane: Function and evolution. J Mol Biol. Academic Press.

21. Mokranjac D, Paschen SA, Kozany C, Prokisch H, Hoppins SC, Nargang FE, Neupert W, Hell K. 2003. Tim50, a novel component of the TIM23 preprotein translocase of mitochondria. EMBO J 22:816–825.

22. Tamura Y, Harada Y, Shiota T, Yamano K, Watanabe K, Yokota M, Yamamoto H, Sesaki H, Endo T. 2009. Tim23 - Tim50 pair coordinates functions of translocators and motor proteins in mitochondrial protein import. J Cell Biol 184:129–141.

23. Singha UK, Hamilton VN, Duncan MR, Weems E, Tripathi MK, Chaudhuri M. 2012. Protein translocase of mitochondrial inner membrane in Trypanosoma brucei. J Biol Chem 287:14480–14493.

24. Duncan MR, Fullerton M, Chaudhuri M. 2013. Tim50 in Trypanosoma brucei possesses a dual specificity phosphatase activity and is critical for mitochondrial protein import. J Biol Chem 288:3184–3197.

25. Singha UK, Hamilton V, Chaudhuri M. 2015. Tim62, a novel mitochondrial protein in Trypanosoma brucei, is essential for assembly and stability of the TbTim17 protein complex. J Biol Chem 290:23226–23239.

26. Harsman A, Oeljeklaus S, Wenger C, Huot JL, Warscheid B, Schneider A. 2016. The non-canonical mitochondrial inner membrane presequence translocase of trypanosomatids contains two essential rhomboid-like proteins. Nat Commun 7.

27. Wenger C, Oeljeklaus S, Warscheid B, Schneider A, Harsman A. 2017. A trypanosomal orthologue of an intermembrane space chaperone has a non-canonical function in biogenesis of the single mitochondrial inner membrane protein translocase. PLoS Pathog 13.

28. Smith JT, Singha UK, Misra S, Chaudhuri M. 2018. Divergent Small Tim Homologues Are Associated with TbTim17 and Critical for the Biogenesis of TbTim17 Protein Complexes in Trypanosoma brucei. mSphere 3.

29. Oelgeschläger T. 2002. Regulation of RNA polymerase II activity by CTD phosphorylation and cell cycle control. J Cell Physiol. John Wiley & Sons, Ltd.

30. Harikrishna Reddy R, Kim H, Noh K, Kim YJ. 2014. The diverse roles of RNA polymerase II C-terminal domain phosphatase SCP1. BMB Rep. The Biochemical Society of the Republic of Korea.

31. Wrighton KH, Willis D, Long J, Liu F, Lin X, Feng XH. 2006. Small C-terminal domain phosphatases dephosphorylate the regulatory linker regions of Smad2 and Smad3 to enhance transforming growth factor-β signaling. J Biol Chem 281:38365–38375.

32. Seifried A, Schultz J, Gohla A. 2013. Human HAD phosphatases: structure, mechanism, and roles in health and disease. FEBS J 280:549–571.

33. Fullerton M, Singha UK, Duncan M, Chaudhuri M. 2015. Down regulation of Tim50 in Trypanosoma brucei increases tolerance to oxidative stress. Mol Biochem Parasitol 199:9–18.

34. Tripathi A, Singha UK, Paromov V, Hill S, Pratap S, Rose K, Chaudhuri M. 2019. The Cross Talk between TbTim50 and PIP39, Two Aspartate-Based Protein Phosphatases, Maintains Cellular Homeostasis in Trypanosoma brucei. mSphere 4.

35. Dawoody Nejad L, Serricchio M, Jelk J, Hemphill A, Bütikofer P. 2018. TbLpn, a key enzyme in lipid droplet formation and phospholipid metabolism, is essential for mitochondrial integrity and growth of *Trypanosoma brucei*. Mol Microbiol 109:105–120.

36. Pelletier M, Frainier AS, Munini DN, Wiemer JM, Karpie AR, Sattora JJ. 2013. Identification of a novel lipin homologue from the parasitic protozoan Trypanosoma brucei. BMC Microbiol 13:101.

37. Kelley, L.A., Mezulis, S., Yates, C.M., Wass, M.N., and Sternberg, M.J.E. (2015). The Phyre2 web portal for protein modeling, prediction and analysis. Nat. Protoc. 10(6),845–858. doi: 10.1038/nprot.2015.053.

38. Trott, O., and Olson, A.J. (2010). AutoDock Vina: improving the speed and accuracy of docking with a new scoring function, efficient optimization, and multithreading. J. Comput. Chem. 31(2),455–461. doi: 10.1002/jcc.21334.

39. Keij JF, Bell-Prince C, Steinkamp JA. 2000. Staining of mitochondrial membranes with 10-nonyl acridine orange MitoFluor Green, and MitoTracker Green is affected by mitochondrial membrane potential altering drugs. Cytometry 39:203–210.

40. Dudek J, Hartmann M, Rehling P. 2019. The role of mitochondrial cardiolipin in heart function and its implication in cardiac disease. Biochim Biophys Acta - Mol Basis Dis. Elsevier B.V.

41. Garcia D, Shaw RJ. 2017. AMPK: Mechanisms of Cellular Energy Sensing and Restoration of Metabolic Balance. Mol Cell. Cell Press.

42. Li Z. 2012. Regulation of the cell division cycle in Trypanosoma brucei. Eukaryot Cell 11:1180–1190.

43. Li Z, Lindsay ME, Motyka SA, Englund PT, Wang CC. 2008. Identification of a bacterial-like HslVU protease in the mitochondria of Trypanosoma brucei and its role in mitochondrial DNA replication. PLoS Pathog 4.

44. Mbang-Benet DE, Sterkers Y, Morelle C, Kebe NM, Crobu L, Portalès P, Coux O, Hernandez JF, Meghamla S, Pagès M, Bastien P. 2014. The bacterial-like HslVU protease complex subunits are involved in the control of different cell cycle events in trypanosomatids. Acta Trop 131:22–31.

45. Sung KH, Lee SY, Song HK. 2013. Structural and biochemical analyses of the eukaryotic heat shock locus v (HslV) from Trypanosoma brucei. J Biol Chem 288:23234–23243.

46. Liu B, Wang J, Yaffe N, Lindsay ME, Zhao Z, Zick A, Shlomai J, Englund PT. 2009. Trypanosomes Have Six Mitochondrial DNA Helicases with One Controlling Kinetoplast Maxicircle Replication. Mol Cell 35:490–501.

47. Serricchio M, Hierro-Yap C, Schädeli D, Ben Hamidane H, Hemphill A, Graumann J, Zíková A, Bütikofer P. 2020. Depletion of cardiolipin induces major changes in energy metabolism in *Trypanosoma brucei* bloodstream forms. FASEB J fj.202001579RR.

48. Meinecke M, Wagner R, Kovermann P, Guiard B, Mick DU, Hutu DP, Voos W, Truscott KN, Chacinska A, Pfanner N, Rehling P. 2006. Tim50 maintains the permeability barrier of the mitochondrial inner membrane. Science (80-) 312:1523–1526.

49. Sugiyama S, Moritoh S, Furukawa Y, Mizuno T, Lim YM, Tsuda L, Nishida Y. 2007. Involvement of the mitochondrial protein translocator component tim50 in growth, cell proliferation, and the modulation of respiration in *Drosophila*. Genetics 176:927–36

50. Malhotra K, Modak A, Nangia S, Daman TH, Gunsel U, Robinson VL, Mokranjac D, May ER, Alder NN. 2017. Cardiolipin mediates membrane and channel interactions of the mitochondrial TIM23 protein import complex receptor Tim50. Sci Adv 3:e1700532.

51. Schnaufer A, Clark-Walker GD, Steinberg AG, Stuart K. 2005. The F1-ATP synthase complex in bloodstream stage trypanosomes has an unusual and essential function. EMBO J. 24:4029–4040

52. Schlame M. 2008. Cardiolipin synthesis for the assembly of bacterial and mitochondrial membranes. J. Lipid. Res. 46:171–199

53. Lilley AC, Major L, Young S, Stark MJR, Smith TK. 2014. The essential roles of cytidine diphosphate-diacylglycerol synthase in bloodstream form trypanosoma brucei. Mol Microbiol 92:453–470.

54. Serricchio M, Butikofer P. 2012. An essential bacterial-type cardiolipin synthase mediates cardiolipin formation in a eukaryote. Proc Natl Acad Sci. E954–961

55. Schädeli D, Serricchio M, Ben Hamidane H, Loffreda A, Hemphill A, Beneke T, Gluenz E, Graumann J, Bütikofer P. 2019. Cardiolipin depletion–induced changes in the *Trypanosoma brucei* proteome. FASEB J 33:13161–13175.

56. Xiao J, Engel JL, Zhang J, Chen MJ, Manning G, Dixon JE. 2011. Structural and functional analysis of PTPMT1, a phosphatase required for cardiolipin synthesis. Proc. Natl. Acad. Sci. USA 108:11860–11865

57. Osman C, Haag M, Wieland FT, Brügger B, Langer T. 2010. A mitochondrial phosphatase required for cardiolipin biosynthesis: the PGP phosphatase Gep4. EMBO J. 29:1976–1987

58. Xu Y, Phoon C., Berno B, D’Souza K, Hoedt E. 2016. Loss of protein association causes cardiolipin degradation in Barth syndrome, Nat. Chem. Biol. 12:641–647.

59. Boynton TO, Shimkets LJ. 2015. Myxococcus CsgA, Drosophila sniffer, and human HSD10 are cardiolipin phospholipases, Genes Dev. 29:1903–1914.

60. Verner Z, Basu S, Benz C, Dixit S, Dobáková E, Faktorová D, Hashimi H, Horáková E, Huang Z, Paris Z, Peña-Diaz P, Ridlon L, Týč J, Wildridge D, Zíková A, Lukeš J. 2015. Malleable Mitochondrion of Trypanosoma brucei. Int Rev Cell Mol Biol 315:73–151.

61. Chaudhuri M, Ott RD, Hill GC. 2006. Trypanosome alternative oxidase: from molecule to function. Trends. Parasitol. 22:484–491

62. Helfert S, Estévez AM, Bakker B, Michels P, Clayton C. 2001. Roles of triosephosphate isomerase and aerobic metabolism in Trypanosoma brucei. Biochem J. 357:117–125

63. Szöőr B, Silvester E, Matthews KR. 2020. A leap into the unknown - early events in African trypanosome transmission Trends Parasitol, 36:266–27

64. Niemi NM, Wilson GM, Overmyer KA, Vögtle FN, Myketin L, Lohman DC, Schueler KL, Attie AD, Meisinger C, Coon JJ, Pagliarini DJ. 2019. Pptc7 is an essential phosphatase for promoting mammalian mitochondrial metabolism and biogenesis. Nat Commun 10:3197

65. Hirumi H, Hirumi K. 1991. In vitro cultivation of *Trypanosoma congolense* bloodstream forms in the absence of feeder cell layers. Parasitology 112:225–36.

