## supplementary material for "*Trypanosoma brucei* Tim50 Plays a Critical Role in Cell Cycle Regulation and Parasite Infectivity"

Running Title: Role of Tim50 in the bloodstream form of *T. brucei*

[a]Department of Microbiology, Immunology, and Physiology, School of Medicine, Meharry Medical College, Nashville, TN, USA

[b]Department of Cell and Developmental Biology, Vanderbilt University, Nashville, TN, USA

*Present address: Minu Chaudhuri, Department of Microbiology, Immunology, and Physiology, Meharry Medical College, Nashville, TN, USA

### Materials and Methods

#### Materials.

All the molecular biology reagents were purchased from Sigma-Aldrich. Custom synthesized oligonucleotides and restriction enzymes were obtained from Thermo Fisher Scientific, USA. Commercially available antibodies were obtained from Thermo Fisher, Sigma-Aldrich and Cell Signaling Company.

#### *T brucei* cell culture and growth rate measurement:

The BF single marker (SM) cell line of *Trypanosoma brucei* 427 was cultured in HMI-9 medium supplemented with 10% fetal bovine serum and 2.5 µg/ml G418 at 37°C in a $CO_2$-incubator (5% saturation) (1). Transfection was conducted using an Amaxa Nucleofector (Lonza, Cologne, Germany) with human T-cell Nucleofector solution and transfectants were selected with 5 µg/ ml hygromycin B (Invitrogen), 2.5 µg/ml phleomycin (Invitrogen) and 10 µg/ml blasticidin (InvivoGen) (2). Cell growth was assessed by inoculating the BF at a cell density of $1 \times 10^5$ cells/ml in fresh medium containing antibiotics in the presence or absence of doxycycline (1 µg/ml). Cells were counted at different growth time points with a Neubauer hemocytometer. The log cumulative cell number was plotted against incubation times.

#### Generation of plasmid constructs and *T. brucei* transgenic cell lines

For in situ tagging, TbTim50 cDNA for the C-terminal fragment (amino acid residues--- ) was PCR amplifed using *T. brucei* genomic DNA as the template and sequence-specific primers (see Table S1 in the supplemental material) and cloned into the pNAT Hygro-12Myc vector (3) to express TbTim50 with a 12 X Myc tag at the C-terminal. The BF(SM) parasites were transfected with the linearized construct and independent clones were obtained by limiting dilution. After selection the positive clones were confirmed by genotyping PCR and used for further analyses. This BF cell line was used as the parental control for our experiments. The TbTim50 RNAi (TbTim50 KD) constructs were generated in p2T7$^{Ti}$-177 RNAi vector as previously described (4) and transfected to the in situ tagged TbTim50-12XMyc BF cells and selected by Phleomycine resistance. In-situ tagged TbHslV-3HA cell line was developed using modified pNAT Hygro-3HA vector as described above.

#### qRT-PCR analysis

For quantitative real-time PCR (qRT-PCR) analysis, RNA was isolated from *T. brucei* cells using RNeasy miniprep isolation kit (Qiagen) and digested with amplification-grade DNase (1 U/µl) (Thermo Fisher) for 1 h. The cDNA was made from 2 µg RNA (DNAse digested) using the iScript cDNA synthesis kit (Bio-Rad). Real-time quantitative PCR was performed with a CFX96 Touch real-time PCR detection system (Bio-Rad). The resulting cDNA was amplified using SsoAdvanced universal SYBR green supermix (Bio-Rad) and gene-specific primers, as indicated (Table S1).

#### Expression of recombinant TbTim50

The entire TbTim50 ORF was PCR amplified by sequence-specific primers (Table S1) and cloned in BamHI and HindIII restriction sites of the pGEX4TI vector for recombinant protein expression. Recombinant proteins were expressed in *Escherichia coli* BL-21 strain. The cultures were grown at 37°C until an optical density at 600 nm of 0.6 to 0.8 was reached and induced with 1 mM isopropyl-β-d-thiogalactopyranoside (IPTG) at 18°C for 16 h. Cells were pelleted by centrifugation at 4,000 rpm for 10 min at 4°C and suspended in 50 mM Tris-HCl (pH 7.5), 2 mM EDTA, 1% Triton X-100, 1 mM dithiothreitol (DTT), and protease inhibitors. Samples were sonicated for 10 cycles, and supernatant was used to affinity purify recombinant protein using glutathione Agarose beads (Thermo Fisher). After incubation with lysate, the glutathione beads were washed with 5 volumes of lysis buffer and the bound proteins were eluted with the Tris-HCl (pH 7.5) buffer containing 10 mm reduced glutathione. Samples were analyzed by Coomassie Brilliant Blue staining to ensure purity of the eluted proteins. The purified proteins were also tested by immunoblot analysis using anti-GST (Thermofisher) antibodies.

**Crude Mitochondria Isolation**

BF cells ($2 \times 10^8$) were resuspended in 500 µl of SMEP buffer (250 mM sucrose, 20 mM morpholinepropanesulfonic acid [MOPS]-KOH, pH 7.4, 2 mM EDTA, 1 mM phenylmethylsulfonyl fluoride [PMSF]) containing 0.03% digitonin and incubated on ice for 5 min. The cell suspension was then centrifuged for 5 min at 6,800 × *g* at 4°C. The resultant pellet was considered the crude mitochondrial fraction, and the supernatant contained soluble cytosolic proteins.

**Phosphatase Assay:**

Phosphatase activity was measured in phosphatase buffer (50-µl reaction mixtures containing Tris acetate, pH 7.4, 1 mm EDTA, 1 mm EGTA, 0.1% BME, and 0.1% ethanol). The substrate for phosphatase activity, p-nitrophenyl phosphate (PNPP), was used at a concentration of 400 mM. Aliquots of 0–60 µg of enzyme (purified recombinant proteins) were preincubated for 1 min at 30 °C. The reaction was initiated by adding a mixture of PNPP at 30 °C. The reactions were quenched by adding 450 µl of 0.25 N sodium hydroxide. Release of p-nitrophenol was determined by measuring the absorbance at A410.

**SDS-PAGE and immunoblot analysis.**

Equal amount of total cellular protein from *T. brucei* were resolved on an SDS-PAGE gel and then transferred to nitrocellulose membranes (Bio-Rad). The antigens were visualized by using the West Pico chemiluminescent substrate (Thermo Fisher). Antibodies against Myc monoclonal antisera (culture supernatant of 9E10 hybridoma clone from ATCC), anti-HA monoclonal antibody (abcam), PIP39 (5), TAO (6), VDAC (7), TbTim17 (8), TbPP5 (9), aldolase (10), and PGK (11) were used as probes. Phospho-AMPK-α (P-Thr172) antibody was purchased from Cell Signaling Technologies. The secondary antibodies used were either anti-rabbit or anti-mouse immunoglobulins linked to horseradish peroxidase (Thermo Fisher).

**Mitochondrial membrane potential:**

Live *T. brucei* cells ($1x10^7$) expressing TbTim50-12X-myc (*in situ*-tagged) were used for MitoTracker® staining as previously described (10). Briefly, MitoTracker® Red CMXROS (Molecular Probe®) in DMSO (1 mM) was added to cells in culture medium to a final concentration of 0.5 µM. The mixture was incubated at 37 ˚C for 10 min. Cells were washed and incubated in fresh culture medium for an additional 30 min. Cells were washed twice with 1x PBS and fixed in 0.37% paraformaldehyde at 4°C for 5 min. Next, cells were centrifuged, washed and resuspended in cold PBS and stored at 4 °C until FACS analysis. Fluorescence intensity (Geom.

Mean) was measured with a FACSCalibur (Becton Dickinson) analytical flow cytometer using absorption at 578 nm and emission at 599 nm.

**Cell cycle analysis by flow cytometry**

TbTim50 RNAi cells (1 x $10^7$) induced with doxycycline for 2 and 4 days and control cells grown for the same time periods were harvested from culture and washed in PBS. Cells were fixed with 150 ul fixative solution (1% triton X-100, 40 mM citric acid, 20 mM sodium phosphate, 200 mM sucrose) and incubate at R.T. for 5 min. After addition of 350 ul of diluent buffer (125 mM $MgCl_2$ in PBS) samples were kept overnight at 4°C. Next 550 µl RNAse solution (preboiled 1.0 mgRNAse/ml in 0.2 M sodium phosphate buffer, pH 7.0) were added and samples were incubate at 37 °C for 3-4 h to remove RNA. After that DNA was stained by addition of propidium iodide at 40 µg/ml and equilibriate at 4°C for several hours in dark. DNA content per cell was measured by flow cytometry (BD FACSCalibur), with excitation at 488 nm. At least 10,000 events were collected for each sample and data were analyzed with FlowJo software.

**Immunofluorescence microscopy**

Cells were stained with MitoTracker as described above and washed twice with PBS. Cells were spread evenly over poly-L-lysine (100 µg/ml in $H_2O$)-coated slides. Once the cells had settled, the slides were washed with cold PBS to remove any unattached cells. The attached cells were fixed with 3.7% paraformaldehyde and permeabilized with 0.1% Triton X-100. After blocking with 5% non-fat milk for 30 min, the slides were washed with 1X PBS. Monoclonal anti-Myc antibody was used as the primary antibody and a fluorescein isothiocyanate (FITC)-conjugated anti-mouse IgG was used as a secondary antibody for visualization under a fluorescent microscope. *T. brucei* aldolase antibody (a generous gift from Meredith Morris, Clemson University) were used to detect aldolae in glycosomes. DNA was stained with 1µg/ml 4′,6-diamidino- 2-phenylindole (DAPI). Cells were imaged using a Nikon TE2000E widefield microscope equipped with a 60× 1.4 NA Plan Apo VC oil immersion objective. Images were captured using a CoolSNAP HQ2 cooled CCD camera and the Nikon Elements Advanced Research software.

**Staining *T. brucei* with Giemsa and DAPI**

For Giemsa-staining, cells were spread onto glass slides, allowed to air-dry at RT and then fixed by methanol 30s. Slides are stained by Giemsa solution ( Sigma). Images were captured in bright field on KEYENCE BZX-xx microscope with 100 X objective. For DAPI-staining cells were spread onto glass slides, allowed to air-dry at RT and then fixed and permeabilised in 100% methanol at −20 °C. Slides were removed from methanol, allowed to air-dry at RT and then rehydrated in 1 × PBS. DAPI (0.1 µg/ml, Sigma) was added on the slides for staining nucleus (N) and kinetoplast (K). Images were taken by KEYENCE BZX-xx microscope. At least 200 cells were examined for each cell type at each time point and the number of cells with 1N1K, 1N2K, 2N2K, and aberrant numbers of NK were counted.

**Multidimensional protein identification technology with label-free relative quantitation of protein levels.**

1x $10^7$ cell of T brucei control and RNAi cell harvested and cell lysates (50 µg proteins) were extracted in radioimmunoprecipitation assay (RIPA) buffer as previous described (12) and immediately processed for trypsin digestion. Prior to LC-MS analysis, protein samples were denatured in 8 M urea and 50 mM Tris-HCl (pH 8.0), reduced with 10 mM Tris(2-

carboxyethyl)phosphine hydrochloride (TCEP) for 60 min, alkylated with 2 mM iodoacetamide for 60 min, and then diluted in 2 M urea with 50 mM Tris-HCl (pH 8.0), at room temperature (RT). Two micrograms of trypsin gold (Promega) was added for overnight digestion (18 h at 37°C), and the tryptic peptides were then immediately desalted using Pierce $C_{18}$ spin columns (Thermo Fischer Scientific) at RT. Peptides were eluted with 80% acetonitrile (ACN) and 0.1% formic acid (FA) and dried completely on a SpeedVac concentrator. For nano-LC-MS/MS analysis, Peptides were resuspended in 5 µl of 0.5% FA and loaded onto a 3-phase MudPIT column (150-µm by 2-cm $C_{18}$ resin, 150-µm by 4-cm strong cation exchange [SCX] resin, filter union, and 100-µm by 12-cm $C_{18}$ resin) as described previously (12). A 10-step MudPIT analysis (0 mM, 25 mM, 50 mM, 100 mM, 150 mM, 200 mM, 300 mM, 500 mM, 750 mM, and 1,000 mM ammonium acetate, with each salt pulse followed by a 120-min acetonitrile gradient of 5 to 50% buffer B [buffer A is 0.1% FA, and buffer B is 0.1% FA in acetonitrile]) was executed for LC-MS analysis using an Eksigent AS-1 autosampler and an Eksigent nano-LC Ultra 2D pump online with an Orbitrap LTQ XL linear ion trap mass spectrometer (Thermo Finnigan) with a nanospray source. MS data acquisition was done by a data-dependent 6-event method (a survey Fourier transform mass spectrometry [FTMS] scan [resolution of 30,000] followed by five data-dependent ion trap [IT] scans for the five consequent most abundant ions). The general mass spectrometric settings were as follows: spray voltage of 2.4 kV, no sheath and no auxiliary gas flow, ion transfer tube temperature of 200°C; collision-induced dissociation (CID) fragmentation (for MS/MS) with 35% normalized collision energy, activation $q$ of 0.25, and activation time of 30 ms. The minimal threshold for the dependent scans was set to 1,000 counts, and a dynamic exclusion list was used with the following settings: repeat count of 1, repeat duration of 30 s, exclusion list size of 500, and exclusion duration of 90 s. For protein identification and quantification, database searches were done with PEAKS 8.5 software against the forward and reverse *T. brucei* and human trypsin sequences (downloaded from GenBank). The parameters for the database search were as follows: full tryptic digestion, up to 3 missed cleavage sites, 20 ppm for peptide mass tolerance, 0.5 Da for fragment mass tolerance, cysteine carbamidomethylation (+57 Da) as a fixed modification, and methionine oxidation (+16 Da) as a variable modification. Relative label-free quantification (LFQ) of the identified proteins was performed with the Q module of PEAKS software based on the extracted ion currents of the identified unique peptides' parent ions or a spectral counting approach, and statistically significant changes were confirmed with Fisher's exact test ($P \leq 0.005$; Benjamini-Hochberg FDR of <0.05).

**Homology modeling and molecular docking**

Protein homology modeling was performed using the Cn3D program available online. Protein structure prediction was performed using the sequence corresponding to the RNA polymerase II CTD phosphatase domain of TbTim50 with the RNA polymerase II CTD phosphatase enzyme (1TA0.pdb) as the starting template by the Phyre2 servers (13). The resulting models were scored, and the quality of the highest scored was validated using various modules in the SAVES server (14). Visualization of structural models were performed using Pymol (Delano). Molecular docking calculations were performed with PA and pNPP structures using Autodock vina (15). All structural biology software was made available through the SBGrid Consortium (16).

**Bioinformatics analysis.**

Bioinformatics pathway enrichment analysis was performed using the database for Annotation, Visualization and Integrated Discovery Bioinformatics resources (DAVID version 6.7) (17), the

Search Tool for the Retrieval of Interacting Genes/Proteins (STRING version 11.0) (18), and the Kyoto Encyclopedia of Genes and Genomes (KEGG) (19) databases and search tools. The identified enriched pathways with *P* values of <0.05 were selected. Significantly enriched pathways were determined using asignificance threshold of an FDR corrected P-value <0.005. Protein family enrichment was done using the STRING database with Protein families (Pfam) as the reference database (20). Functional domain analysis of proteins was performed using InterPro version 84.0 to classify protein families and predicted active domains (21).

**Electron microscopy.** Cells were fixed in 2% (v/v) glutaraldehyde, and 2% (w/v) paraformaldehyde in o.1 M sodium cacodylate buffer (SCB), pH 7.2. Cells were then washed with SCB, post-fixed with 1% (w/v) osmium tetroxide in SCB, stained with 0.5% aqueous magnesium uranyl acetate, dehydrated, and embedded in Spurr's resin (22). Blocks were sectioned at 50-70 nm thickness and stained with 5% (w/v) uranyl acetate in 1% acetic acid, and 0.4% lead citrate in 0.1 N NaOH. Section grids were inserted into FEI CM12 twin lense 420 transmission electron microscope to capture images.

**Densitometry and statistical analyses.** ImageJ software (NIH) was used to perform densitometry of Western blots. The band intensity of the loading control was used for normalization, and values were plotted using MS Excel. Standard errors were calculated from three independent experiments for each blot. The data were compared with the values for the control and analyzed for statistical significance by using the PRISM (GraphPad) *t*-test. *P* values are indicated by asterisks in the figures.

**Table S1.** Top ranked poses of the theoretical binding free energies evaluated through the AutoDock Vina scoring function

| SUBSTRATE | COMPUTED BINDING FREE ENERGY (ΔG; KCAL/MOL) |
|---|---|
| Phosphatidic Acid (PA) | -4.8 |
| para-Nitrophenyl Phosphate (pNPP) | -4.7 |

*Putative binding site residues were identified as those interacting with the substrate molecule within 3.0Å (using ChimeraX) and were the same for both substrates: ***D242, D244, E245, S250, T299, A300, Y336, D354, N355, S356***. PA's phosphate group interacts with Y336 while the phosphate group of pNPP interacts with D242.

**Table S2.** Primers used in this study

| Name | Sequence (5'---3') | Length (bp) |
|---|---|---|
| TbTim50RNAi F | AGTCGGATCCGCATAGAGGGGAAAAGAGTGAGG | 33 |
| TbTim50RNAi R | AGTCAAGCTTGGACGAAAAGCAACATAAACGGTG | 33 |
| TbTim50-Myc P1 | GATAGGATCCATGAACCACGATGCCATGTC | 30 |
| TbTim50 (1269)- P2 | GATCCTCGAGTCAGTAATGCGGAGAGTTTTGTC | 33 |
| TbTim50-(703) P3 | GGTCAAGCTTACAAGATCACACTTATATTAGATC | 34 |
| TbTim50-pNAT P4 | GGATCCTCACCTAGGCAGGTCTTCTTCAGA | 30 |
| TbHslV-HA P1 | GATCAAGCTTTGTTGTCATTGGTTTCGCGG | 30 |
| TbHslV-HA P2 | GATCTCTAGACTCGCTAGTTTTCGCCGAC | 29 |
| rGST-TbTim50 F | GATAGGATCCATGAACCACGATGCCATGTC | 30 |
| rGST-TbTim50 R | GATCCTCGAGTCAGTAATGCGGAGAGTTTTGTC | 33 |
| EP1- qRT-PCR F | GAAGGACCAGAAGACAAGGG | 20 |
| EP1- qRT-PCR R | AGGTTCAGGCTCAACTTCGT | 20 |
| TbPAD2-qRT-PCR F | GGTCTCGCCTTCCCAGCATT | 20 |
| TbPAD2-qRT-PCR R | TCAGCGTTATCGTCGCAGGT | 20 |
| TbTim50-qRT-PCR F | CCGCCTCCGTCTCGGTTTAT | 20 |
| TbTim50-qRT-PCR R | CCAGGTCCCGACCAAGCAAT | 20 |
| TERTqRT-PCR F | GAGCGTGTGACTTCCGAAGG | 20 |
| TERTqRT-PCR R | AGGAACTGTCACGGAGTTTGC | 21 |
| PIP39qRT-PCR F | TGGAGGTGCAGGTGTTACAA | 20 |
| PIP39qRT-PCR | CGTTTGAGTTCGGGAAAGGG | 20 |

**Figure S1**

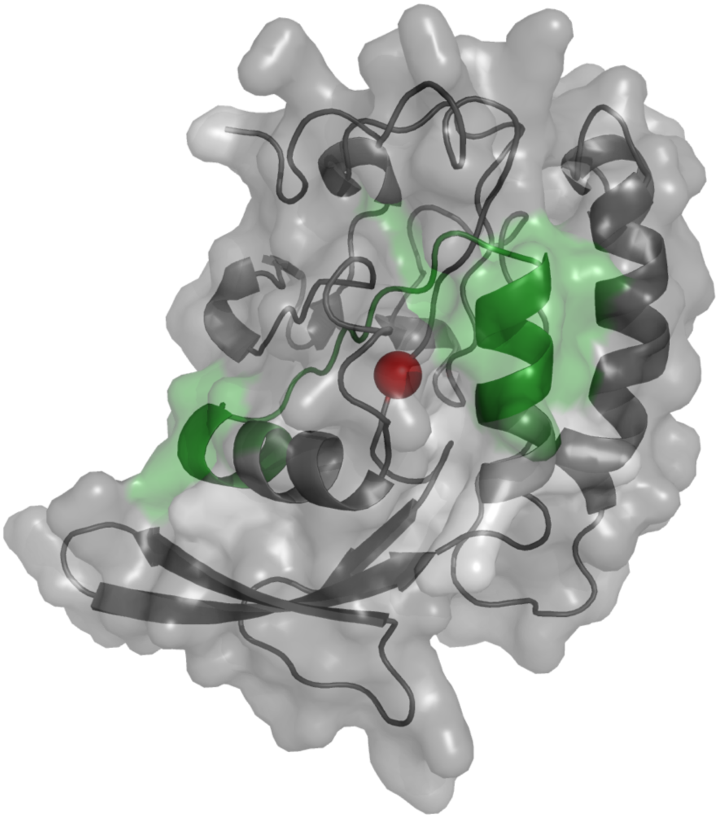

**FIG. S1.** Structural homology model of functional TbTim50. The structural homology model of the CTD PPase domain of TbTim50 (228-404 amino acids) highlighting the proximity of the active site D242 (red sphere) and the location of the predicted transmembrane segment (285-304 amino acids) shown as green helices.

Figure S2.

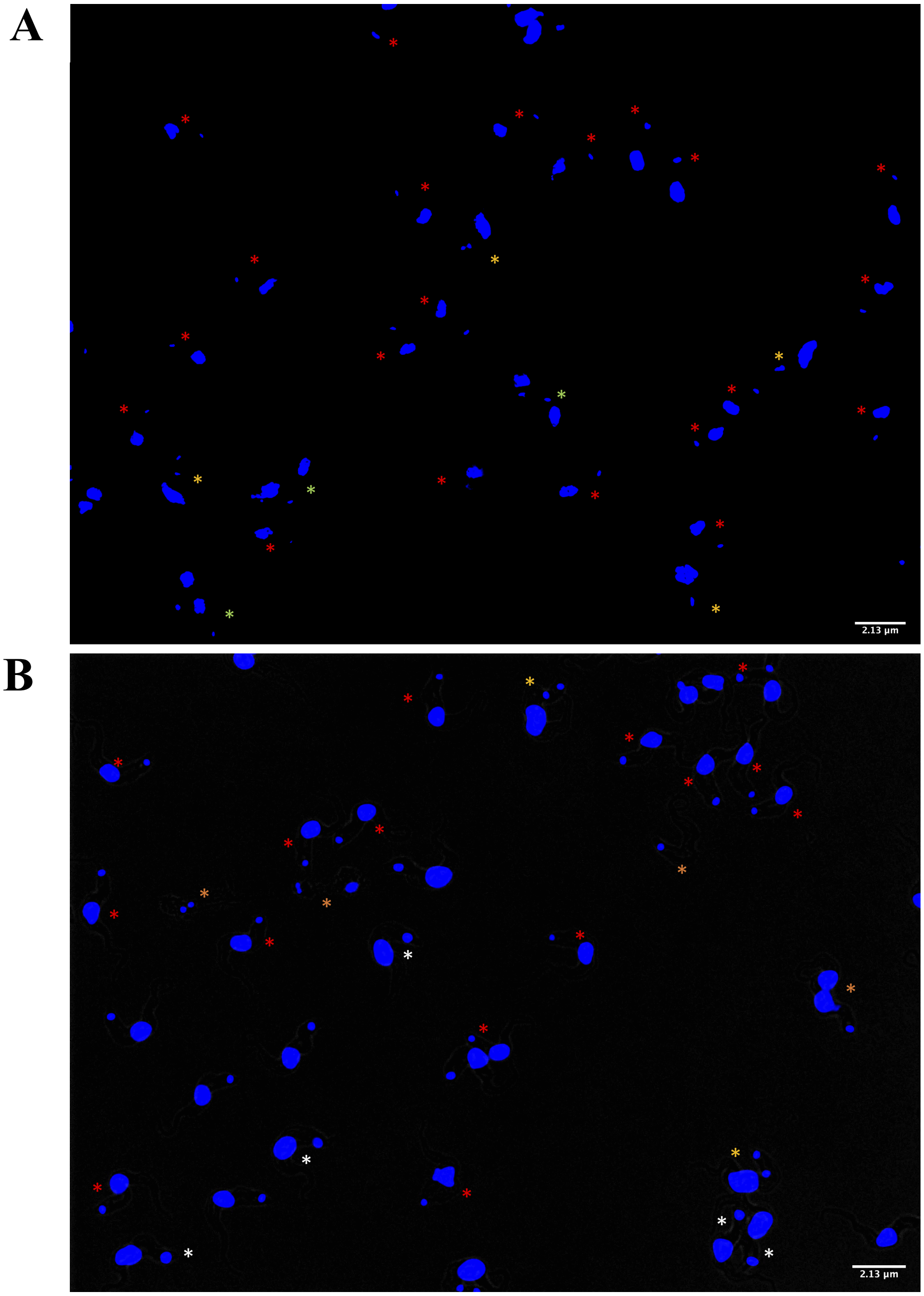


**FIG. S2.** Fluorescence microscopy of DAPI-stained BF before (A) and after (B) induction of TbTim50 RNAi for 4 days. Cells containing 1K1N, 2K1N, 2K2N, larger KN are marked with red, yellow, green and white asterisks.

Figure S3

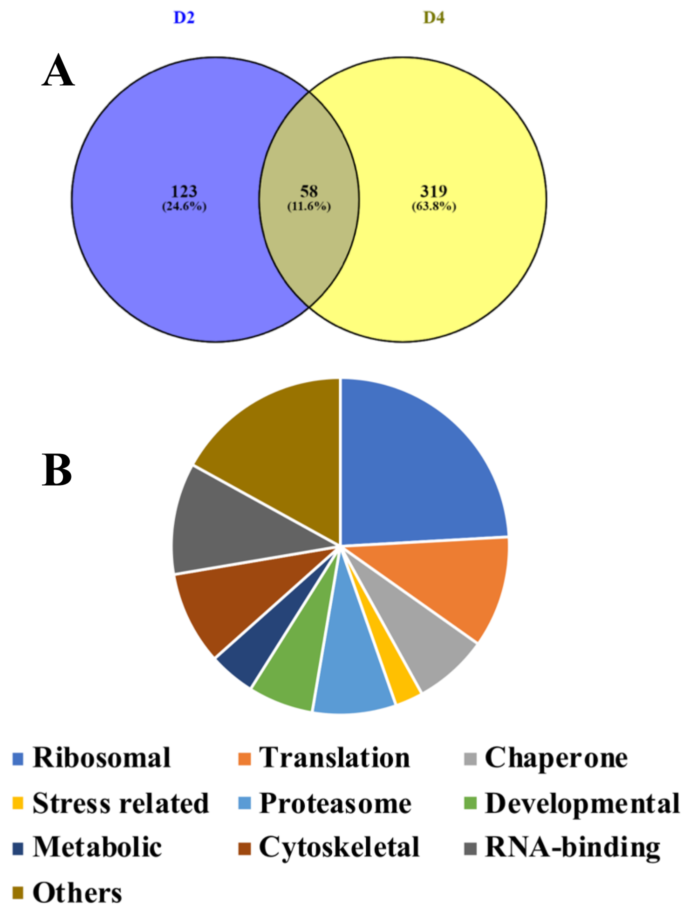


**FIG. S3.** Changes in BF proteomes due to TbTim50 KD. (A) Semi-quantitative proteomics analyses were performed using control and TbTim50 RNAi cells grown for 2 and 4 days in the presence of doxycycline. Overlap of significantly altered proteins identified 58 common proteins. The levels of these proteins were changed in cells at 2 (D2) and 4 (D4) days after RNAi. B. The common proteins were classified according to their functional terms and presented in a pi-chart.

### References for supplement


1. Hirumi H, Hirumi K. 1991. In vitro cultivation of *Trypanosoma congolense* bloodstream forms in the absence of feeder cell layers. Parasitology 112:225–36.
2. Amaxa
3. Alsford
4. Duncan MR, Fullerton M, Chaudhuri M. 2013. Tim50 in *Trypanosoma brucei* possesses a dual specificity phosphatase activity and is critical for mitochondrial protein import. J Biol Chem 288:3184–3197.
5. Szöor B, Ruberto I, Burchmore R, Matthews KR. 2010. A novel phosphatase cascade regulates differentiation in *Trypanosoma brucei* via a glycosomal signaling pathway. Genes Dev 24:1306–1316.
6. Chaudhuri M, Ajayi W, Hill GC. 1998. Biochemical and molecular properties of the *Trypanosoma brucei* alternative oxidase. Mol Biochem Parasitol 95:53–68.
7. Singha UK, Sharma S, Chaudhuri M. 2009. Downregulation of mitochondrial porin inhibits cell growth and alters respiratory phenotype in *Trypanosoma brucei.* Eukaryotic Cell 8:1418–28.
8. Singha UK, Peprah E, Williams S, Walker R, Saha L, Chaudhuri M. 2008. Characterization of the mitochondrial inner membrane translocator Tim17 from *Trypanosoma brucei*. Mol Biochem Parasitol 159:30-43.
9. Chaudhuri M. 2001. Cloning and characterization of a novel serine/threonine protein phosphatase type 5 from *Trypanosoma brucei*. Gene 266:1–13
10. Misset O, Bos OJ, Opperdoes FR. 1986. Glycolytic enzymes of *Trypanosoma brucei*. Simultaneous purification, intraglycosomal concentrations and physical properties. Eur J Biochem 157:441-5.
11. Parker HL, Hill T, Alexander K, Murphy NB, Fish WR, Parsons M. 1995. Three genes and two isoenzymes: gene conversion and the compartmentalization and expression of the phosphoglycerate kinases of *Trypanosoma* (Nannomonas) *congolense*. Mol Biochem Parasitol. 69:269-279
12. Lin, AJ, Eng J, Schieltz DM, Carmack E, Mize GJ, Morris DR, Garvik BM, Yates JR 3rd. 1999. Direct analysis of protein complexes using mass spectrometry. Nat Biotechnol. 17: 676–682.
13. Kelley, LA, Mezulis S, Yates CM, Wass MN, and Sternberg MJE. 2015. The Phyre2 web portal for protein modeling, prediction and analysis. *Nat. Protoc.* 10: 845-858
14. Benkert P, Kunzli M., Schwede T. 2009. QMEAN server for protein model quality estimation. Nuc. Acid Res. 37:W510-W514
15. Trott O, and Olson AJ. 2010. AutoDock Vina: improving the speed and accuracy of docking with a new scoring function, efficient optimization, and multithreading. J. Comput. Chem. 31:455-461
16. Morin A, Eisenbraun B, Key J, Sanschagrin PC, Timony MA, Ottaviano M, Sliz P. 2013. Cutting Edge: Collaboration gets the most out of software. eLife 2:e01456
17. Huang DW, Sherman BT, Lempicki RA. Systematic and integrative analysis of large gene lists using DAVID Bioinformatics Resources. *Nature Protoc. 2009;4(1):44-57.* STRING
18. Szklarczyk D, Gable AL, Lyon D, Junge A, Wyder S, Huerta-Cepas J, Simonovic M, Doncheva NT, Morris JH, Bork P, Jensen LJ, von Mering C.

STRING v11: protein-protein association networks with increased coverage, supporting functional discovery in genome-wide experimental datasets. Nucleic Acids Res. 2019 Jan; 47:D607-613

19. Kanehisa M, Furumichi M, Tanabe M, Sato Y, Morishima K. KEGG: new perspectives on genomes, pathways, diseases and drugs. Nucleic Acids Res. 2017 Jan 4;45(D1):D353-D361. doi: 10.1093/nar/gkw1092. Epub 2016 Nov
20. Huang DW, Sherman BT, Lempicki RA. Bioinformatics enrichment tools: paths toward the comprehensive functional analysis of large gene lists. Nucleic Acids Res. 2009;37(1):1-13**.**
21. Blum M, Chang H, Chuguransky S, Grego T, Kandasaamy S, Mitchell A, Nuka G, Paysan-Lafosse T, Qureshi M, Raj S, RichardsonL, Salazar GA, Williams L, Bork P, Bridge A, Gough J, Haft DH, Letunic I, Marchler-Bauer A, Mi H, Natale DA, Necci M, Orengo CA, Pandurangan AP, Rivoire C, Sigrist CJA, Sillitoe I, Thanki N, Thomas PD, Tosatto SCE, Wu CH, Bateman A and Finn RD The InterPro protein families and domains database: 20 years on. *Nucleic Acids Research, Nov 2020*EM
